# Markers of resilience to stony coral tissue loss disease and probiotic potential in the microbiome of the threatened coral, *Orbicella faveolata*

**DOI:** 10.64898/2025.12.21.695810

**Authors:** Cynthia C. Becker, Allison R. Cauvin, Karen L. Neely, Caroline Dennison, Andrew C. Baker, Brian K. Walker, Julie L. Meyer

## Abstract

Stony coral tissue loss disease (SCTLD) is widespread within the Caribbean and affects at least 22 species of reef-building coral. Bacteria have been implicated in the etiology of SCTLD, but the community of bacteria and archaea may also contribute to SCTLD resistance. To identify potential mechanisms through which microbes contribute to SCTLD resistance, we sequenced metagenomes from 41 colonies of the threatened coral, *Orbicella faveolata,* in the lower Florida Keys. All colonies were fate-tracked for three to five years and disease lesions were treated with amoxicillin. By 2024, 20% were never diseased, 10% had lesions before sampling but recovered, 22% were apparently healthy but were eventually susceptible to infection, and 49% had regular repeated infections. Within the coral microbiome, diseased and yet-to-be diseased colonies exhibited higher variability in functional genes. In contrast, corals that remained unaffected or recovered had less variable microbiomes with greater abundances of vitamin and antibiotic biosynthesis, secretion system, and quorum sensing genes that may support host health and resilience to pathogens. Though on some colonies antibiotic treatments were applied repeatedly, there was no effect on the diversity of beta-lactamases, antibiotic resistance genes that may confer amoxicillin resistance. Additional potentially probiotic gene clusters for the production of antimicrobial and bioactive compounds were present in many colonies regardless of fate. Taken together, we find significant probiotic potential in the coral microbiome to armor host *O. faveolata* corals against SCTLD infection, which may underpin intraspecific variation in stony coral tissue loss disease resilience and susceptibility.

## Introduction

Stony coral tissue loss disease (SCTLD) is a contagious disease affecting at least 22 species of reef-building corals in Florida and the Caribbean [1]. The outbreak began in Florida in 2014, and resulted in significant losses to coral colony density and coral cover [2, 3], near extirpation of some highly susceptible species [4], and decline in ecosystem functions such as reef accretion [5]. While some corals are highly susceptible to this disease, others show varying susceptibility and resist infection for years [reviewed by 1, 6]. The mountainous star coral, *Orbicella faveolata,* which is listed as threatened under the U.S. Endangered Species Act and as endangered by the IUCN, is a bouldering, reef-building coral historically common in the Caribbean with moderate susceptibility to SCTLD relative to other corals [7]. Within a diseased reef, some *O. faveolata* colonies seem resistant and do not develop lesions even over years of exposure [8], while others nearby quickly and repeatedly develop numerous lesions [9, 10]. The underlying mechanisms contributing to SCTLD resistance by some colonies remains an open area of research.

Resistance to SCTLD may be due to any number of coral traits within the holobiont. The coral holobiont consists of the coral host and its symbiotic dinoflagellate algae (Symbiodiniaceae), bacteria, archaea, viruses, and fungi [11]. Here, we focus on the role of the microbial communities within the holobiont and their potential for supporting host disease resistance and susceptibility. We define the coral microbiome as the prokaryotic component of the holobiont (bacteria, archaea), which is known to support coral host health as well as cause disease [12, 13]. This microbiome confers benefits to the holobiont by cycling nitrogen [14], providing essential vitamins [15], and providing probiotic benefits like antibiotic defenses [16, 17]. However, the microbiome may also cause diseases [18, 19]. While the causative agent of SCTLD remains elusive, the lesions harbor potentially pathogenic genera [20–25]. Some colonies even have co-infections with *Vibrio coralliilyticus* bacteria [26]. Therefore, investigating the microbiome within the host offers a promising avenue to better understand SCTLD resistance.

While 16S rRNA gene sequencing provides insight into bacteria that may play a role in the disease, this approach lacks information on functional genes that would inform disease etiology. For example, many studies associate *Vibrio* with SCTLD [20, 21, 24, 26], but healthy coral hosts also often harbor *Vibrio* bacteria which play a beneficial role in nitrogen acquisition [27]. Additionally, 16S rRNA gene sequencing is often unable to differentiate *Vibrio* species, and therefore may misidentify *Vibrio* marine pathogens [28, 29]. For this reason, we focused on metagenomics-based approaches, which have previously shed light on SCTLD pathogenesis [30–32]. Extending this previous research, we apply metagenomics-based methods to identify microbial markers that may shape host resistance to infection.

Incorporating symbiont composition into resistance analyses is important for understanding SCTLD dynamics. In addition to the bacterial communities, zooxanthellate scleractinian corals form an obligate symbiosis with photosynthetic dinoflagellates in the Family Symbiodiniaceae [33, 34]. Although the host relies on the symbionts, the algal symbionts are independently susceptible to disease that can affect their mutualism with the coral host. In the case of SCTLD, histopathology and field observations suggest involvement of the Symbiodiniaceae in affected corals [35]. It is hypothesized that disruption (and eventual degradation) of the Symbiodiniaceae is the first sign of disease which eventually leads to SCTLD lesions [35]. In situ observations support this idea because SCTLD quiesces upon coral paling or bleaching, a sign of the coral host expelling its algal symbionts [9, 36–38]. Along with Symbiodiniaceae density in the coral host, the species of symbiont may affect SCTLD susceptibility [39].

To identify microbial signatures of disease resistance, we sequenced metagenomes from apparently healthy tissue of 41 fate-tracked *O. faveolata* coral colonies in the lower Florida Keys (Figure 1, Table S1). These corals had known health histories before and after sampling, and those with SCTLD lesions were treated with topical amoxicillin paste to preserve tissue and prevent mortality [9, 10, 40]. To identify markers of disease resilience within the coral microbiome, we compared corals of distinct SCTLD fates.

**Figure 1.**
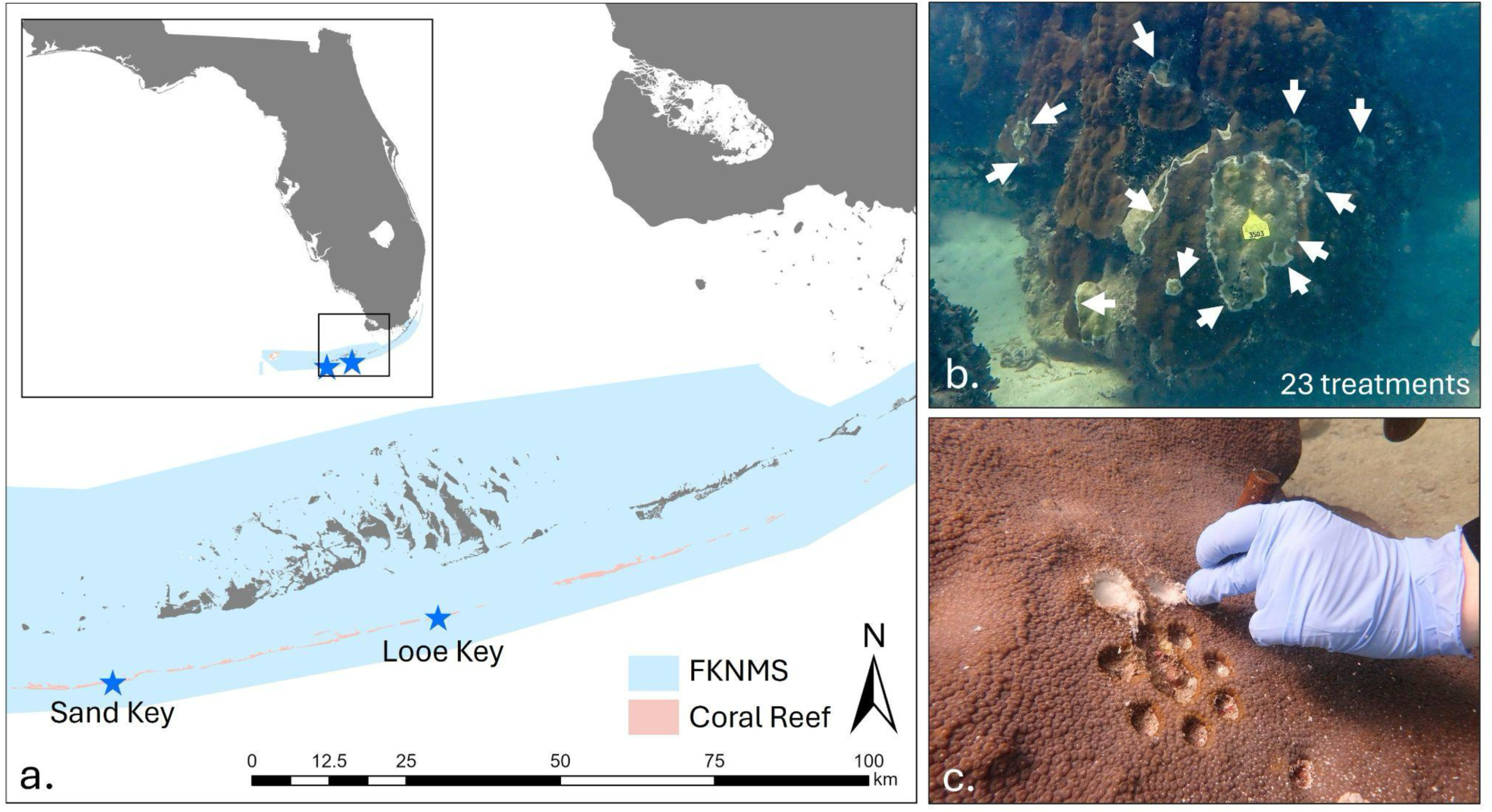
a) Location of the two sampling sites (Sand Key and Looe Key) within the lower Florida Keys and the Florida Keys National Marine Sanctuary (FKNMS). b) Example of an *Orbicella faveolata* with numerous amoxicillin treatments (arrows; 23 were recorded for this coral on this date, but not all are visible in this photo). Note that treatments are highly variable in size and are also spatially dispersed across the colony, with some tissue isolates not having any treatments and some having many. c) Collection of sample from an *O. faveolata* colony using a 12 mm leather punch. Adjacent cores were collected for other collaborative projects.

Additionally, given the variable susceptibility of colonies to SCTLD and the sometimes repeated treatment with amoxicillin, we investigated the presence of diverse antibiotic resistance genes and other biosynthetic gene clusters that produce antimicrobial compounds and their associations to antibiotic treatments.

## Methods

### Study area and collection

Core samples were taken from 41 *Orbicella faveolata* colonies at two reef sites in the lower Florida Keys: Looe Key and Sand Key (Figure 1a). The selected corals were part of a long-term SCTLD intervention and monitoring project across multiple species and sites [9, 10, 40]. Samples from the selected corals were collected by the SCTLD Resistance Research Consortium (RRC) to measure over a dozen holobiont traits [41]. In-water amoxicillin interventions and fate-tracking were initiated at Looe Key in March 2019 and at Sand Key in October 2020. Each site was visited approximately every two months through July 2024, and any coral affected by SCTLD was tagged with a numbered cattle tag, mapped and photographed for revisitation, and treated on active disease lesions with a topical amoxicillin paste (See SI Methods). All previously tagged corals were monitored for health status during each visit. Based on the health histories of these corals, *O. faveolata* at both sites were selected for further analysis as part of the RRC [41]. The selected colonies were intentionally chosen to represent a range of susceptibility levels.

Coral cores were collected via SCUBA on 12-13 June 2021. Separate 12 mm leather punches were used to extract a 1 cm core from apparently healthy tissue on each colony far from active SCTLD lesions (Figure 1C). Tissue cores were placed into a labeled bag. At the surface, they were preserved in DNA/RNA Shield (Zymo Research, Irvine, CA, USA) and kept on ice. All samples were stored at −80°C until processing.

### Colony characteristics

Following over three years of fate-tracking, we developed a category of colony fate that incorporated the full health histories of the colonies. Each colony was classified as either: 1) “diseased” (had SCTLD before sampling, during sampling, and after sampling), 2) “will be diseased” (was healthy at the time of sampling, but developed SCTLD lesions at some point afterwards – time of lesion development varied from 4 – 30 months after sampling), 3) “recovered” (coral had SCTLD before sampling, but not during sampling and at no point after sampling), or 4) “unaffected” (coral was never observed with SCTLD). Collaborative partners assessed coral host genotypes and algal symbiont composition for each colony (See SI Methods).

### DNA Extraction and Library preparation

For extraction of prokaryotic DNA, coral cores were thawed on ice, then 1-2 polyps of coral tissue and mucus were scraped from cores using flame-sterilized scalpels and forceps. The tissue was placed in 1mL of phosphate-buffered saline with ethylenediaminetetraacetic acid and mechanically broken up with a sterile micropestle. The resulting slurry was added to the Qiagen QIAamp DNA Microbiome Kit (Qiagen, Germantown, MD, USA), and DNA was extracted according to the manufacturer’s protocol. Four reagent blanks were generated alongside experimental samples, without coral biomass. The extracted DNA was then submitted to the University of Florida’s Interdisciplinary Center for Biotechnology Research NextGen DNA Sequencing core (RRID:SCR_019152) for library preparation and sequencing on an Illumina NovaSeq 6000 (Illumina, San Diego, CA, USA) in 150-bp paired-end format. Further details are in the SI Methods.

### Bioinformatics

An overview of the bioinformatics pipeline we employed to target bacteria and archaea within the coral holobiont is outlined in Figure S1. Detailed parameters can be found in the SI Methods. We assessed the quality of raw reads using FastQC v.0.11.7, then filtered the raw reads using trimmomatic v.0.39 [42]. To remove host and endosymbiont DNA and begin enriching for prokaryotic genomic content, we used Bowtie2 v.2.4.2 [43] to map trimmed reads against the following genomes: *O. faveolata*, *Breviolum minutum*, *Symbiodinium* sp. Clade C, and *Durusdinium sp.* We assembled the remaining reads that did not map to the eukaryotic genomes (host/endosymbiont-cleaned reads) from each of the 41 corals individually using MegaHit v.1.1.3 and assessed the assemblies with QUAST v.5.2.0 [44] (Table S2). These assemblies are available at Zenodo DOI 10.5281/zenodo.11493775 [45]. To evaluate the broad prokaryotic versus eukaryotic content of the assemblies, we used EukRep v.0.6.7 [46] followed by QUAST v.5.2.0.

Following taxonomic assignment of contigs, we further enriched the prokaryotic component of the host/endosymbiont-cleaned reads (Table S2). We mapped the reads to the eukaryotic portion of the metagenome assembly (from EukRep) to generate eukaryote-removed reads with Bowtie2 v.2.4.2 [43]. We then generated a co-assembly of all 41 coral samples with MegaHit v.1.1.3 [47], then evaluated it with QUAST v.5.2.0. We used EukRep v.0.6.7 again to keep the prokaryotic portion of the co-assembly for annotation and analysis.

To predict protein-coding genes, we used Prodigal v.2.6.3 [48]. We annotated the predicted genes (amino acid fasta) with MMseqs2 v.2/14, employing the GTDB database v.214.1 [49]. For functional annotations, we used GhostKOALA [50]. We quantified the abundance of predicted genes from the prokaryote co-assembly using Salmon v.1.10.1 [51] to generate a table of read counts per sample for each predicted gene in the prokaryote co-assembly. The predicted proteins, annotations, and abundance table are publicly available at Zenodo 10.5281/zenodo.17947166 [45].

We used the resistance gene identifier (RGI) v.6.0.2 to find antibiotic resistance genes (ARGs) in the predicted genes from the prokaryote co-assembly (amino acid fasta). To identify biosynthetic gene clusters (BGCs), we used antiSMASH v.7.0.0 on the prokaryotic co-assembly generated by EukRep. We manually curated the antiSMASH and MIBiG results to identify antimicrobial BGCs (Table S3). CoverM v.0.6.1 was used to measure mean coverage of BGCs in the eukaryote-removed reads [52] that was converted to presence/absence for statistical analysis.

### Statistical analyses

For statistical tests, we used the read counts for each coral sample of the predicted genes in the coassembly (247,254 genes). We analyzed the beta diversity of functional genes using Aitchison distance to account for compositional data [53]. To test how colony-specific covariates (coral genotype, fate, disease condition at sampling,

Symbiodiniaceae composition, and reef location) explained variability in the functional microbiome, we used the adonis2 function [54] in vegan v.2.6.6.1. For all tests, genotypic clones were treated as separate individuals (Figure S2a). To investigate dispersion of the prokaryotic microbiome we used the betadisper function in vegan v.2.6.6.1. Then, we conducted a Kruskal-Wallis or ANOVA test followed by either a Wilcoxon or Tukey post-hoc test.

To identify genes that may be involved in disease resistance, we conducted a differential abundance test in corncob between unaffected corals (baseline) and the other colony fates. We used corncob v.0.4.1 [55] only on the 31,695 functionally annotated genes and controlled for the composition of Symbiodiniaceae. Significant differences were evaluated using the parametric Wald test, with a Benjamini-Hochberg false discovery rate correction of 0.05. Prior to plotting, KEGG annotations were retrieved for each KO ID using the KEGG API in R [56].

To test whether repeated amoxicillin treatments impacted the richness and diversity of beta-lactamase ARGs, we used the package breakaway v.4.8.4 [57] to estimate the richness of beta-lactamase genes. We tested for the effect of amoxicillin treatments and recent treatment history on richness using the betta function [58]. We additionally tested for the effect of these variables on the beta diversity of beta-lactamase genes using the methods described above. To test whether BGCs were differentially prevalent across colony fates, we used a Fisher’s Exact Test incorporating a Monte-Carlo simulation, followed by a Benjamini-Hochberg multiple test correction in R. Further statistical detail is in the SI Methods.

## Results

### Disease history and characteristics of *Orbicella faveolata* colonies

*Orbicella faveolata* colonies from two reefs in the lower Florida Keys were monitored for up to two years prior to sampling in June 2021 for stony coral tissue loss disease (SCTLD) and treated in cases of active lesions (Figure 1). The number of total topical amoxicillin treatments applied to colonies before sampling ranged from 0 (on colonies that had never been observed with SCTLD) to 779 (Table S1). The colonies with the most treatments were generally large (up to 32 m^2^ of estimated live tissue area), and often had new lesions during each visit (Figure 1B; Tables S1).

After three to five years of bimonthly observations, 20% of the selected colonies were never observed with SCTLD (unaffected). In contrast, 49% exhibited SCTLD lesions before, during, and after sampling (diseased), 10% of colonies were diseased prior to sampling but resisted infection for the entire post-sampling period (recovered), and 22% were ultimately susceptible to disease even if they were unaffected during sampling (will be diseased) (Figure 2). During tissue sampling, 49% exhibited active disease lesions and 51% were apparently healthy.

**Figure 2.**
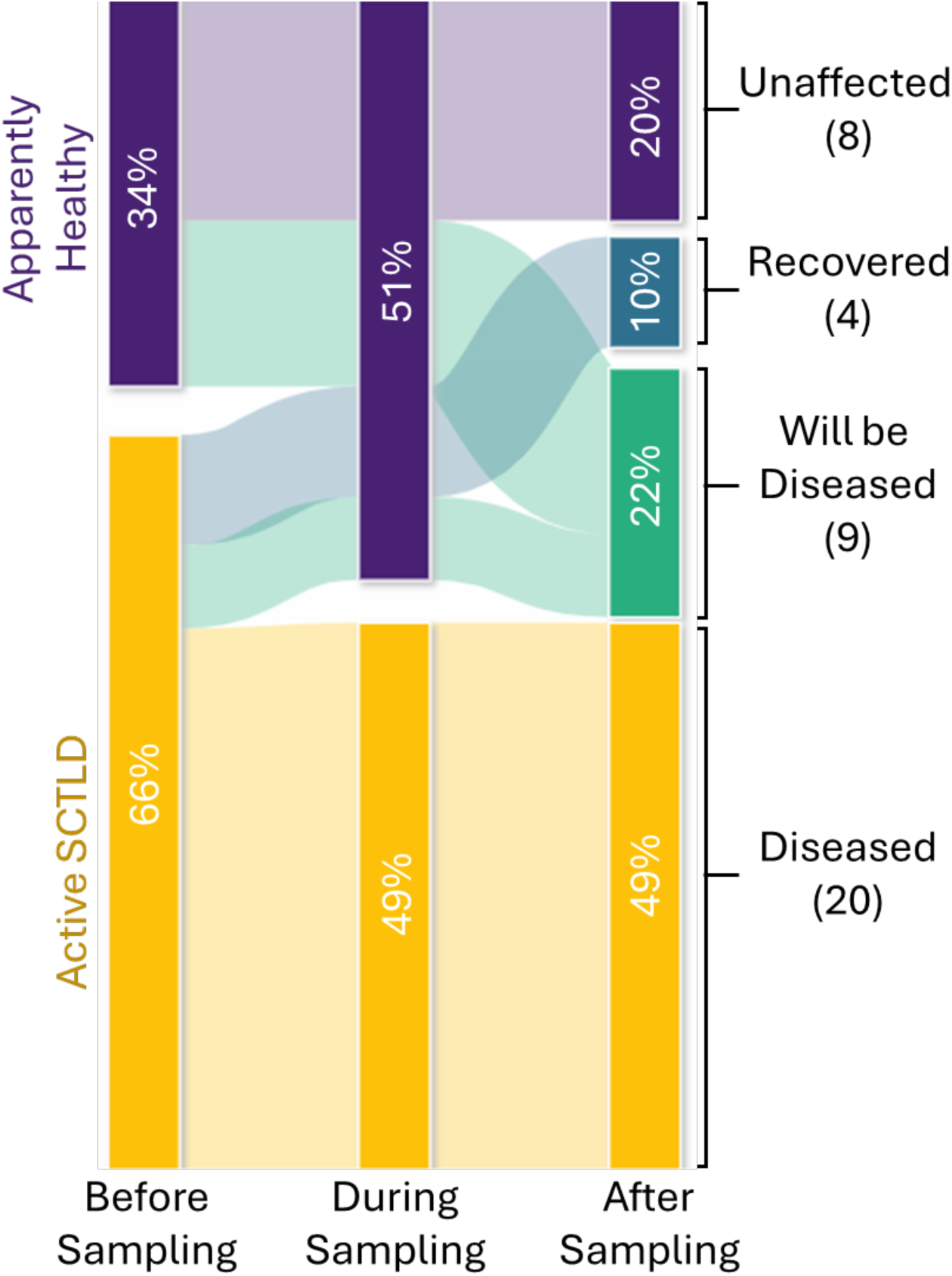
We fate-tracked 41 *Orbicella faveolata* colonies every two months for up to five years for the presence of stony coral tissue loss disease. The presence of SCTLD at any point before sampling (as early as 2019), during sampling (June 12-13, 2021), or after sampling (July 2021 - July 2024) defined four “colony fate” categories: unaffected, recovered, will be diseased, and diseased. The number of colonies in each category are represented in parentheses.

We quantified Symbiodiniaceae community composition, and for all but one coral sample, *Breviolum* was the main genus of Symbiodiniaceae present (Figure S3). 46.3% of the colonies harbored small proportions of *Cladocopium, Durusdinium,* or both genera in addition to *Breviolum.* One coral sample (3421) contained both *Cladocopium* and *Durusdinium* genera, but no *Breviolum*.

### *In silico* enrichment of the coral prokaryotic microbiome

Sequencing of apparently healthy coral holobiont metagenomes resulted in approximately 132 million reads per sample that were filtered to approximately 30.4 million reads per coral sample after selecting for prokaryotic genomic content (Table S2). In addition, we generated 16S rRNA analyses of four DNA extraction reagent blanks (See SI Results). We removed on average 74.35% of reads deemed eukaryotic (See Methods, Table S2). This left, on average, 26% of reads per sample that were combined into a co-assembly. The prokaryotic portion of the co-assembly was 11% of the total length of the assembly, was 287,437,943 bp long, had an N50 of 1,903, and a longest contig of 367,269 bp. Following protein-coding gene prediction, the co-assembly contained 247,254 genes that were used for downstream analyses.

### Markers of disease resilience in the coral functional microbiome

Symbiodiniaceae community composition, colony fate, and colony condition at sampling explained 53.4% of the variation in coral functional microbiome beta diversity (PERMANOVA, 999 permutations, p < 0.05, Figure 3, Table S4). Symbiodiniaceae community composition and colony fate explained 32.8% and 12.7% of the variance respectively (Figure 3, Table S4). Condition of the colony at sampling (apparently healthy or actively diseased) explained 7.9% of the variance (Figure 3, Table S4). Reef and coral host genotype did not explain microbiome composition (PERMANOVA p > 0.05, Figure S2).

**Figure 3.**
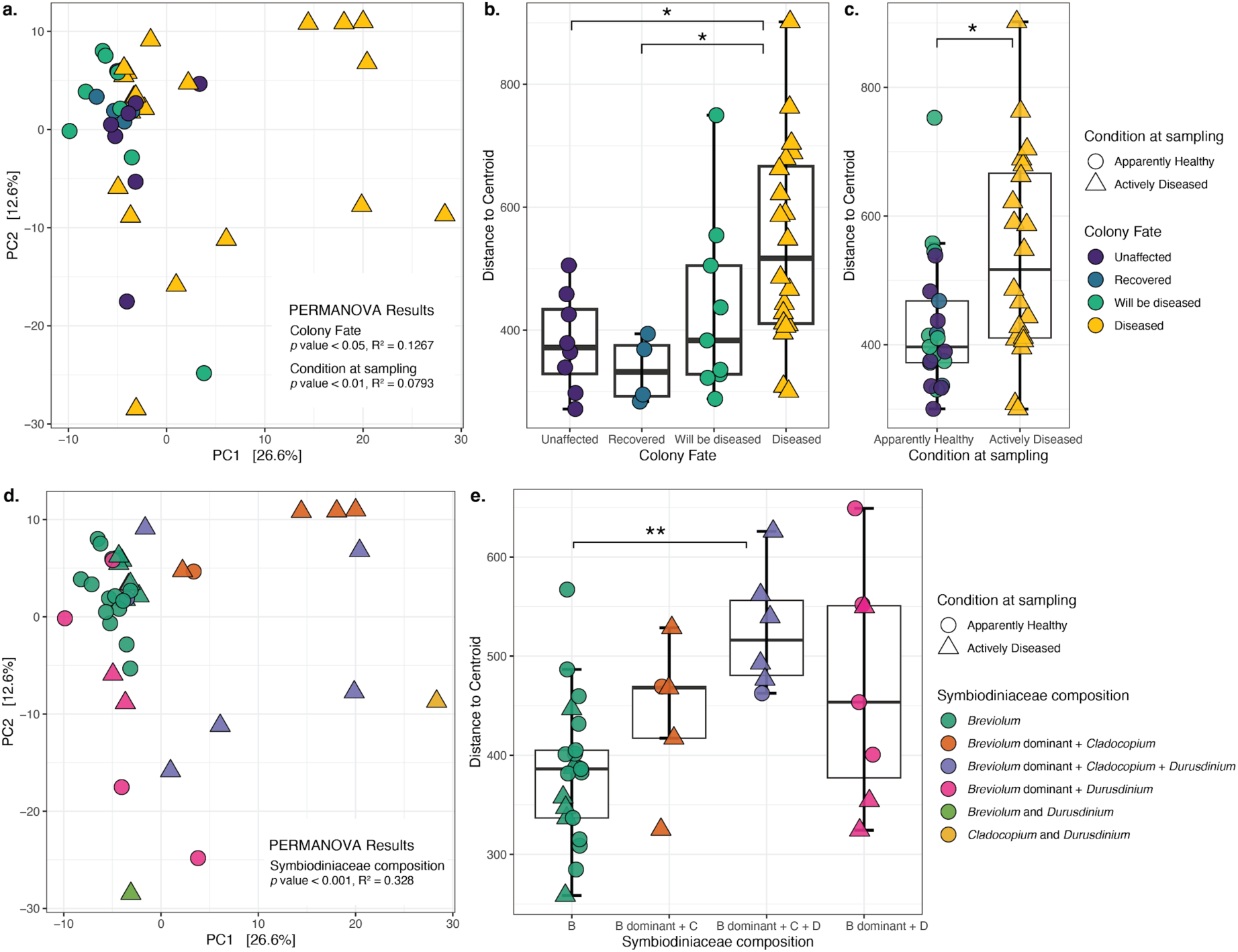
Functional microbiome composition in *Orbicella faveolata* is explained by colony fate, coral condition at sampling, and Symbiodiniaceae composition (PERMANOVA, 999 permutations, p < 0.05). a) Principal components analysis (PCA) of coral tissue prokaryotic functional microbiome beta diversity (Aitchison distance) colored by colony fate with point shape reflecting the condition of the coral at sampling. b) Prokaryotic functional beta diversity dispersion measured as the distance to centroid was greater in colonies with diseased fates relative to recovered or unaffected fates (Wilcoxon rank sum test, *p < 0.05). c) Prokaryotic functional beta diversity dispersion measured as the distance to centroid was greater in actively diseased colonies relative to apparently healthy colonies (Wilcoxon rank sum test, *p < 0.05). d) PCA as in (a) colored by Symbiodiniaceae composition. e) Functional prokaryotic beta dispersion differed across *Symbiodiniaceae* composition groups, with corals containing *Breviolum* dominant + *Cladocopium* + *Durusdinium* exhibiting greater dispersion than *Breviolum-*only tissues (Tukey Honest Significant Differences post-hoc test, **p < 0.01). Note that the *Cladocopium* and *Durusdinium* and the *Breviolum* and *Durusdinium* groupings are omitted from panel (d) because there was only one coral in each group. All points are colored by *Symbiodiniaceae* composition groupings, and shape reflects the colony condition at the time of sampling. The groupings *Breviolum* and *Durusdinium,* and *Cladocopium* and *Durusdinium* contain both genera making up at least 10% of symbiont community. See Figure S3 for a visualization of the photoendosymbiont composition of all corals.

Dispersion in the coral prokaryotic functional microbiome was increased in actively diseased colonies relative to apparently healthy colonies (Wilcoxon rank sum test p < 0.05, Figure 3c). Functional microbiomes from diseased coral fates were more dispersed compared to those of recovered and unaffected fates, but not to those of the will be diseased fate (Wilcoxon Rank Sum post-hoc test, adjusted p < 0.05, Figure 3b). For corals in the will be diseased fate, the microbial dispersion was not related to months until disease onset, nor to presence or absence of disease lesions prior to sampling (linear model or Wilcoxon rank sum test, p > 0.05, Figure S4). Increased functional microbiome variability also coincided with differences in Symbiodiniaceae composition (ANOVA p < 0.001, Figure 3e). Compared to colonies with exclusively *Breviolum* symbionts, those with small portions of both *Cladocopium* and *Durusdinium* harbored more dispersed prokaryotic functional microbiomes (Tukey HSD post-hoc test adjusted p < 0.05, Figure 3e).

We identified 392 functional genes that were differentially abundant in the recovered, will be diseased, and diseased coral fates relative to corals unaffected for the duration of our monitoring period (Corncob adjusted p < 0.05, Table S5). Colonies that recovered from a prior SCTLD infection and remained disease-free during the duration of the study (recovered) harbored greater abundances of vitamin and cofactor metabolism genes, quorum sensing genes, beta-lactam resistance genes, membrane transport genes such as secretion systems, and other signalling genes (Figure 4, Table S5). In contrast, unaffected tissue of actively diseased colonies had lower overall abundances of most genes relative to the genes in unaffected colonies, including vitamin metabolism genes and secondary metabolite biosynthesis genes encoding multiple antibiotics (phenazine, novobiocin, carbapenem, monobactam), membrane transport genes, quorum sensing, and beta-lactam resistance genes (Figure 4, Table S5). Despite the colonies being apparently healthy at sampling, corals with will be diseased fates had microbiomes with lower abundances of most cofactor and vitamin metabolism genes, secondary metabolite biosynthesis genes, and transport and signalling genes relative to unaffected colony fates, which mirrored the patterns of diseased colonies (Figure 4, Table S5).

**Figure 4.**
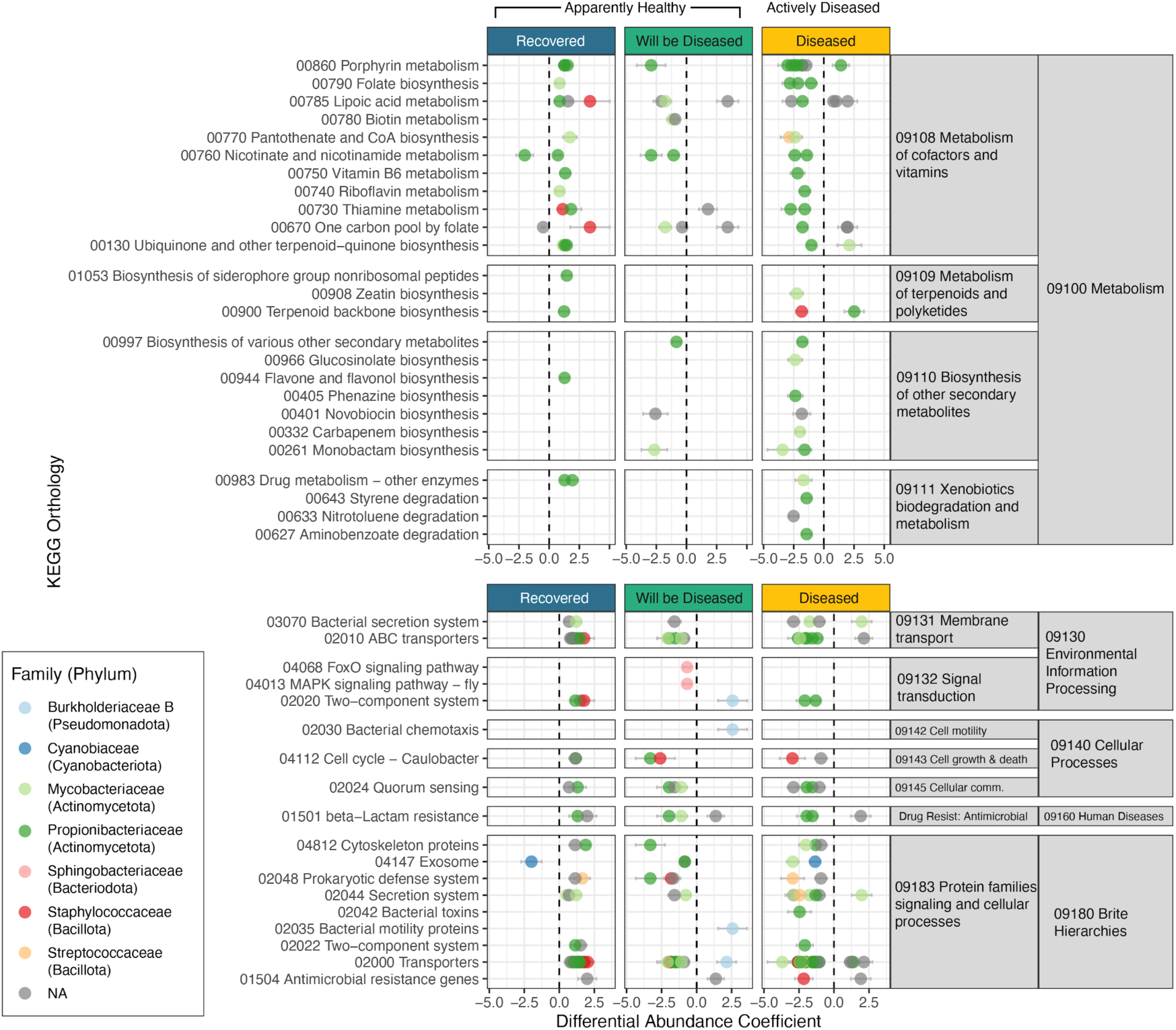
Genes involved in metabolism of secondary metabolites and other natural products, as well as those involved in signal transduction and cellular processes were less abundant in diseased and corals that will be diseased categories as compared to unaffected colonies, but enriched in corals recovered from SCTLD (Corncob, Benjamini-Hochberg false discovery rate corrected *p* < 0.05). The differential abundance coefficient represents how much the modeled relative abundance of a gene was enriched or depleted relative to the baseline (unaffected colonies, dotted line at zero). Error bars reflect 95% confidence intervals around the coefficient. Results are organized by KEGG hierarchy, with the top panel representing metabolism genes, and the bottom panel representing all other processes. There are some points (genes) that are repeated due to their involvement in multiple pathways and roles. The color represents the Family and Phylum-level annotations for a given predicted gene from the GTDB database. These differentially abundant genes are a subset of all. See Table S6 for a list of all 392 genes and the comparisons for which they were significantly differentially abundant.

### Coral microbiomes encode diverse probiotic potential for antimicrobial resistance and biosynthesis

The 14 colonies that were never-diseased prior to sampling were never treated with amoxicillin, while the other 27 with previous SCTLD lesions were treated at some point (and sometimes repeatedly) before sampling (Table S1). Despite the potential selective pressure exerted by repeated application of up to hundreds of treatments applied over the two years prior to metagenomic sampling, we did not find a correlation between the number of amoxicillin treatments (raw and scaled to m^2^ of live coral tissue) or treatment history (categorical: never treated, treatment > 2 months prior, recent treatment < 2 months prior) and the richness of beta-lactamase genes within the coral microbiome (betta test p > 0.05, Figure 5a). Beta diversity of beta-lactamase genes also did not differ due to number of treatments or treatment history (PERMANOVA p > 0.05, Figure 5b). Of the 53,467 beta-lactamase genes present in the coral microbiome, most were taxonomically unclassified, but some were classified as *Actinomycetia*, *Alphaproteobacteria*, *Clostridia*, or *Gammaproteobacteria* (Figure 5c, Table S6).

**Figure 5.**
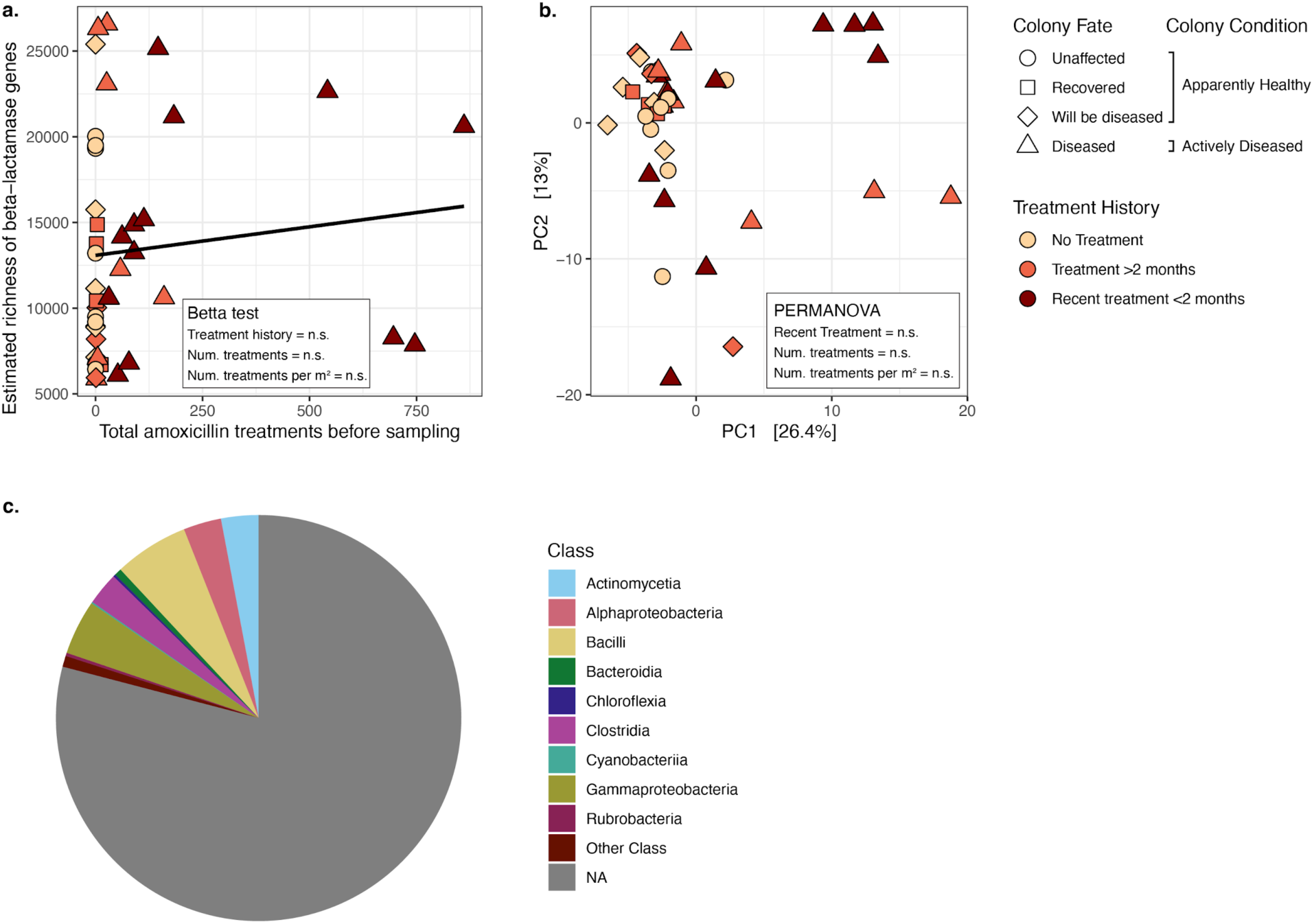
Diverse beta-lactamase genes exist within the coral microbiome regardless of application of the antibiotic amoxicillin. a) The estimated richness (breakaway) of the beta-lactamase genes in each sample did not significantly increase with amoxicillin treatments or recent amoxicillin application (betta test p > 0.05 for total number of amoxicillin treatments (continuous), total number of amoxicillin treatments scaled per m^2^ of live tissue area (continuous), and treatment history (categorical)). The line represents a linear regression. n.s. = not significant from the betta test for heterogeneity of total diversity. b) Principal component analysis showing the beta diversity of beta-lactamase genes in the coral microbiome did not differ due to number of treatments (raw and scaled to tissue area) or recent treatment application (PERMANOVA, p > 0.05). Points in a) and b) are shaped based on the colony fate and colored based on treatment history. c) Taxonomic affiliation of the 53,467 individual beta-lactamase genes depicted as a pie chart indicated most were unclassified (NA), and a small proportion were annotated at the class taxonomic level. The top nine most abundant classes are shown and the “Other Class” includes the remaining 94 classes.

We investigated the diverse biosynthetic gene clusters (BGCs) within the coral microbiome and identified 102 BGCs, of which 56 encoded putatively probiotic antimicrobial compounds (Figure 6, Table S3, Figure S5). The 56 BGCs encoding antimicrobial compounds were prevalent across coral colonies, regardless of their colony fate (Fisher’s Exact Test, Benjamini-Hochberg adjusted p > 0.05, Figure 6). They included BGCs that matched biosynthetic genes for a variety of marine-derived compounds, such as curacin A, bryostatin, and polytheonamide A and polytheonamide B (Figure 6). Some of the most highly prevalent BGCs had matches to xenematide, toyoncin, ergovaline, carnobacteriocin XY, strobilurin A, and polytheonamide A and B (Figure 6). Overall, the closest matches in the MIBiG database to the coral microbiome BGCs identified in this study encoded 65 different compounds, of which 43 were antimicrobial or toxins (Table S3, Figure S5).

**Figure 6.**
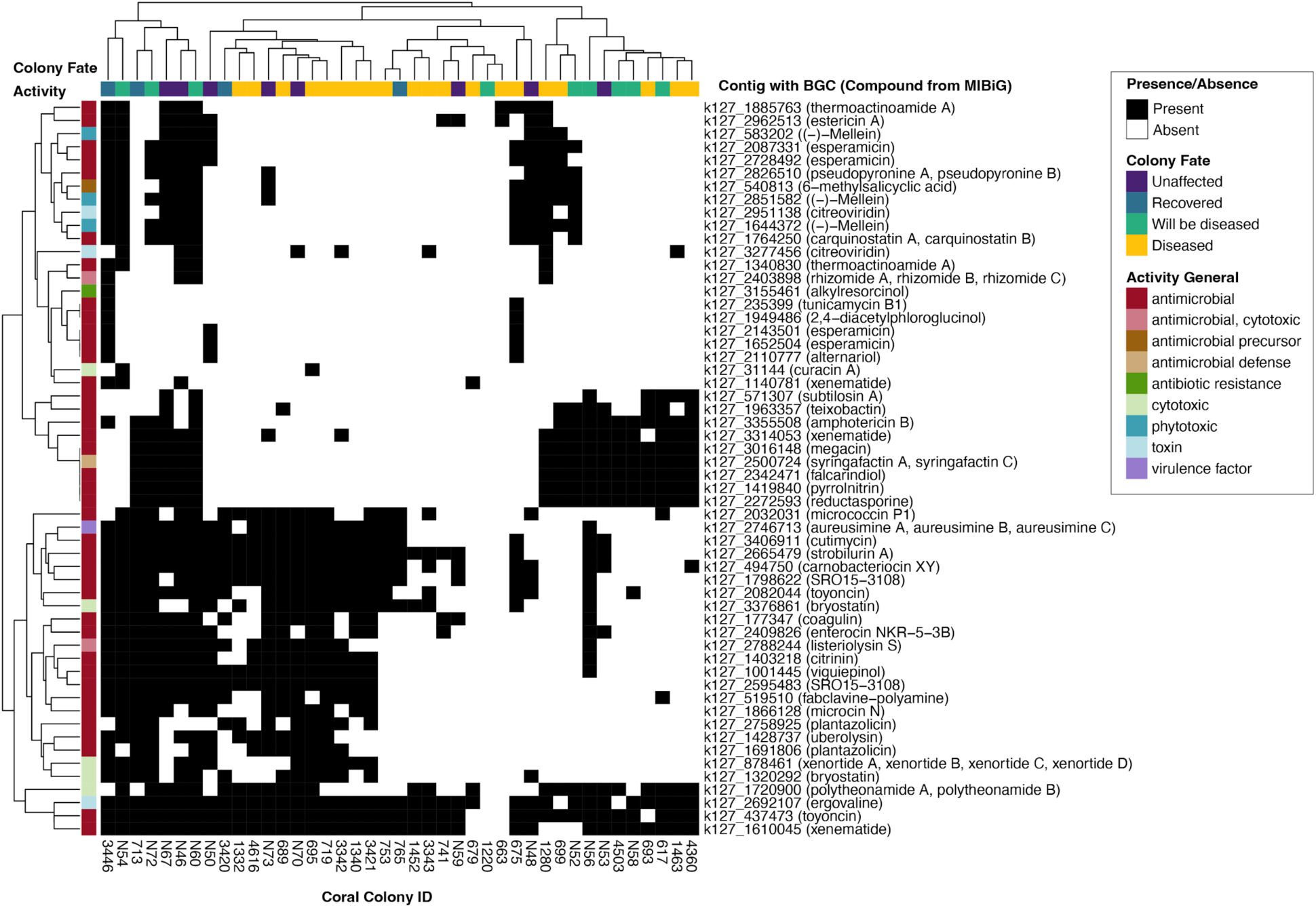
The coral microbiome encodes diverse potential for antimicrobial activity regardless of disease condition (Fisher’s Exact Test, Benjamini-Hochberg false discovery rate corrected p > 0.05). Heatmap displays presence or absence of a contig containing the potential antimicrobial-encoding biosynthetic gene cluster (BGC) across all coral colonies and is clustered by Euclidean distance. Presence was defined as a non-zero mean coverage of the contig in the eukaryote-removed reads and the compound label reflects the compound encoded by the closest MIBiG database match returned by antiSMASH (see methods). Coral colonies are colored to reflect the fate of the colony and contigs are colored to reflect the inferred activity of the BGC.

## Discussion

We undertook 3.5 years of bimonthly fate-tracking and antibiotic interventions on *O. faveolata* colonies and identified intraspecific variation in resistance to SCTLD. Following deep sequencing of the prokaryotic functional microbiome, we found evidence that vitamin biosynthesis and probiotic genes are associated with resilience, and may improve host health in the face of disease exposure. Conversely, dysbiotic functional microbiomes were a marker of disease. In these large *O. faveolata* colonies, the microbiome harbored diverse beta-lactamase genes, even in colonies that were never treated with amoxicillin. As stony coral tissue loss disease continues to impact reefs with no known etiological agent or fully effective cure, these data provide insight into the microbial markers that may underlie the variable susceptibility of *O. faveolata* colonies to SCTLD.

Within apparently healthy tissue of *O. faveolata,* colonies impacted by SCTLD had more variable microbial communities than unaffected individuals, which may be an indicator of systemic dysbiosis. The Anna Karenina hypothesis offers a useful analogy for dysbiosis in animal microbiomes that are diseased: “All happy families are alike; each unhappy family is unhappy in its own way” [59]. In our study, all recovered and unaffected colonies harbored low microbial variability, while diseased colonies were more dispersed, reflecting dysbiosis. Interestingly, will be diseased colonies, which were apparently healthy during sampling but became diseased following sampling, had intermediate microbial functional variability. Although the increased variability within the will be diseased colonies did not relate to time until infection or to past infection history, this overall stochasticity of the microbiome within healthy corals prone to infection may be a marker of SCTLD. Previous 16S rRNA taxonomic investigations found increased dispersion associated with healthy tissue in SCTLD-affected colonies [21, 24], and here we identify microbial functional dispersion in corals not yet infected with SCTLD. Therefore, SCTLD may first present as a functional dysbiosis within the coral microbiome that opens it up to disease [32]. SCTLD-susceptible corals are known to be dysbiotic over time [60], and while we did not sample multiple timepoints for metagenomics here, we hypothesize that those disease-susceptible *O. faveolata* colonies would be most variable in their functional microbiome. Contrasting this, in previous coral bleaching work, lower microbial functional dispersion was associated with increased probiotic potential [61]. Here, our recovered and unaffected coral colonies were those with the lowest dispersion. In contrast to a dysbiosis signal, our work suggests resilient and resistant corals are not just those with specific microbial taxa, but those armored with a consistent suite of microbial functions.

Functional stochasticity in the coral microbiome also coincided with varied Symbiodiniaceae composition. Corals with only *Breviolum* symbionts generally had less microbial dispersion than coral colonies that also included small amounts of *Cladocopium* and *Durusdinium*. Connection between Symbiodiniaceae, a large component of the coral holobiont, with the prokaryotic microbiome has been reported previously [62, 63]. Additionally, the Symbiodiniaceae-associated microbiome can influence the cnidarian host fitness [64]. Here, the composition of Symbiodiniaceae explained 33% of prokaryotic functional variability, linking algal symbionts to the coral microbiome. Further, we suggest that bacterial-based resilience may additionally be linked to the underlying algal community in SCTLD-susceptible corals.

Microorganisms within resilient, or recovered, corals harbored increased potential to support host health compared to corals that ultimately succumbed to disease.

Recovered coral microbiomes had higher relative abundances of B vitamin metabolism genes. Corals are unable to produce B vitamins, and coral-associated bacteria likely transfer them to coral and symbiont hosts to support their metabolism [15, 65].

Additionally, quorum-sensing was a marker of resilient corals. While quorum sensing is associated with coral disease and bleaching [66, 67], some of the differentially abundant quorum sensing genes here included components of DAHP synthase and Anthranilate synthase, important in tryptophan amino acid biosynthesis [68]. Tryptophan is exuded by corals [69], and non-pathogenic coral-associated bacteria exhibit chemotaxis toward it [70]. Here, quorum sensing capabilities may be important for maintaining natural coral-microbial symbioses and increasing host resilience to disease. In contrast to recovered corals, actively diseased and apparently healthy corals that contracted disease after sampling had microbiomes with lower abundances of vitamin B metabolism and quorum sensing genes, which may have compromised host fitness. Additionally, compromised coral microbiomes had lower abundances of secondary metabolite and xenobiotics biosynthesis pathways. These pathways can be important for defense against infection [71], and lowered potential for these may have additionally predisposed these colonies to the disease they ultimately contracted.

Antibiotic application with the beta-lactam amoxicillin is the primary intervention strategy for halting SCTLD lesions in the Caribbean [6, 9, 40, 72–74]. Although effective at preventing SCTLD-related mortality, these treatments have raised concerns over the potential development of antibiotic resistance [75, 76]. There was no evidence in our study that treatments selected for an increased richness or diversity of beta-lactamase genes, even with corals that received hundreds of individual lesion treatments up to two years prior to sampling. Our findings were complicated by the variability in treatment history of each coral. One treatment was defined as the application of amoxicillin paste onto an active lesion, regardless of lesion size, location on the colony relative to the sample location, or colony size. For example, some of the largest colonies received hundreds of treatments that were often meters from the sampling location. Additionally, these applications occurred one month to over two years prior to sampling. We accounted for these varied complications by scaling treatments to colony size and testing if treatment history impacted beta-lactamase diversity. Our findings offer an optimistic view that repeated topical antibiotic treatments did not select for greater potential for antibiotic resistance within the coral microbiome. Additionally, recent findings indicate these localized, topical treatments do not affect coral physiology [77], reproduction [78, 79], or gene expression [80], and treatments do not appear to increase the expression of antibiotic resistance genes (ARGs) [81]. Indeed, we identified about 200,000 ARGs, representing the majority of the 247,254 protein-coding prokaryotic genes in the co-assembly. Similarly, ARGs were prevalent in untreated, SCTLD-impacted coral tissue [31]. Our research indicates that antibiotic resistance may not be a signal of disease treatments, but rather a native feature of the coral microbiome [81]. While encouraging, antibiotics are known to select for antibiotic resistance, and moving forward, managers and scientists aiming to curb SCTLD should continue to be prudent with environmental applications of amoxicillin.

Antimicrobial and bioactive biosynthetic gene clusters were prevalent across coral microbiomes, regardless of colony fate, and may play an important role in host defense. We identified 102 biosynthetic gene clusters (BGCs) within the prokaryotic functional microbiome, and of these, 56 encoded potential antimicrobial or toxin production genes (Figure 5, Figure S5). The detection of broad and diverse pathways for antimicrobial production may not be surprising given the known ability of coral-associated bacteria to synthesize antimicrobial compounds that inhibit potential pathogens [82–84]. For example, *Pseudoalteromonas* bacterial isolates from healthy *O. faveolata* in SCTLD-impacted areas are known to produce the antibiotic alteramide A [85]. Considering specific *Pseudoalteromonas* strains are a promising probiotic for treating SCTLD [16, 86], investigating the suite of other antimicrobial products may shed light into the native ways corals may resist SCTLD infection, without the addition of probiotics. While we didn’t specifically detect genes for the synthesis of alteramide A, our in silico approaches identified multiple other antibiotic and bioactive production pathways. Only three matched natural products isolated and described from marine environments, highlighting the paucity of marine research compared to those in terrestrial, agricultural, or human disease-related environments. These compounds: curacin A, bryostatin, and polytheonamide A and B, all play a role in host defense and have demonstrated anti-cancer activity. Curacin A was isolated from the marine cyanobacterium, *Lyngbya majuscula,* from Caribbean reef habitats and kills both human cell lines and brine shrimp [87]. We found genes encoding production of this compound present in one apparently healthy and one diseased *O. faveolata.* Bryostatins may provide a protective role for host larvae by deterring predation [88]. Like curacin A, bryostatin has also demonstrated anticancer and cytotoxic activity [89]. Two bryostatin biosynthetic gene clusters identified in both actively diseased and apparently healthy *O. faveolata* microbiomes match (0.11 - 0.16 similarity score) those from the uncultured symbiont of a marine bryozoan, *Candidatus* Endobugula sertula [90]. Polytheonamide A and polytheonamide B were isolated from a Pacific reef sponge, *Theonella swinhoei,* and show cytotoxic activity against cancer cells [91]. These products are likely synthesized by uncultivated *Candidatus* Entotheonella bacterial symbionts [92], which are known to harbor immense bioactive potential, and may serve as chemical defense for the sessile host sponge [93]. Like bryostatin and curacin A, genes for polytheonamides A and B were present in metagenomes of both apparently healthy and actively diseased colonies. Considering we sampled apparently healthy tissue on all colonies, perhaps the presence of genes encoding production of cytotoxic compounds is a reflection of the native and significant potential for coral microbiomes to harbor bacteria that protect the coral host.

Some of the most prevalent biosynthetic gene clusters were those that may play a role in host defense and homeostasis. Xenematide, toyoncin, and ergovaline production genes were present in over 90% of the corals, and carnobacteriocin XY and strobilurin A were present in 68% of the sampled corals. Xenematide is produced by bacterial symbionts of nematodes to maintain symbiosis of the microbe within the nematode host [94]. Toyoncin is a bacteriocin isolated from a *Bacillus cereus* strain that is antimicrobial against foodborne pathogens [95]. While mainly considered a food contaminant, *B. cereus* has also been isolated from marine sponges, where it also produces antibiotics [96]. Ergovaline and strobilurin A are both fungal-based antimicrobial compounds [97, 98], though the nearest BLAST-based matches to the core biosynthetic gene from antiSMASH were bacterial, suggesting similar antimicrobial compounds across domains or potential horizontal gene transfer events. Carnobacteriocin XY was isolated from lactic acid bacteria, which are common in food, though *Carnobacterium* species that produce antimicrobial compounds are also found in marine habitats [99]. The high prevalence of these antimicrobial and defensive compounds is likely reflective of the *O. faveolata* microbiome’s ability to support host homeostasis and prevent predation or infection, which would be essential within apparently healthy tissue, regardless of infection status of the coral host overall.

While some *O. faveolata* colonies exhibited repeated SCTLD infections, over three to five years of in situ monitoring, 20% of our intentionally selected corals resisted infection, and 10% were resilient and recovered from SCTLD. By combining this rich disease history with the characterization of the functional potential of bacterial microbiomes, we provide evidence that this resilience to disease may be due to the ability of the coral microbiome to synthesize antimicrobial products that protect against invading pathogens. The survival of these large coral colonies was also due to antibiotic interventions that halted disease progression, yet these treatments did not alter the natively diverse suite of antibiotic resistance genes present within the coral microbiome. In addition to identifying markers of resilience, we uncovered high microbial dispersion, or dysbiosis, present in unaffected colonies that may have predisposed them to the disease they contracted. While causative agents of this widespread disease remain elusive, our efforts uncovered a suite of probiotic markers of resilience to SCTLD within the coral microbiome. This supports the growing emphasis on developing probiotic treatments and microbial consortia to improve coral host fitness in the face of disease, and other widespread stressors [16, 100].

## Data Availability Statement

The raw sequence reads are publicly available in the NCBI Sequence Read Archive under BioProject accession number PRJNA925892. The individual coral metagenome assemblies, the co-assembly, predicted proteins, annotations, and abundances are publicly available via Zenodo dataset https://doi.org/10.5281/zenodo.11493775 version 2.0. Scripts containing the code with parameters for bioinformatics as well as the code to reproduce the analyses and figures within this manuscript are available on GitHub at https://github.com/meyermicrobiolab/FLK_SCTLD_Resistance_OFAV/.

## Acknowledgements

This research was made possible by the Florida Department of Environmental Protection Office of Resilience and Coastal Protection, Southeast Region (award numbers B8A48D, C0CB08, C2205D). Coral monitoring was partially supported by funding to KLN in 2019-2023 through Florida Department of Environmental Protection Coral Protection and Restoration Program (awards B40346, B54DC0, B77D91, B967BC, and C01957) and the National Fish and Wildlife Foundation in 2022-2023 (award 0302.21.071754). Colony tagging, treating, and monitoring were conducted under Florida Keys National Marine Sanctuary permits 2018-141, 2019-115, and 2020-077. Samples were collected under Florida Keys National Marine Sanctuary permit FKNMS-2021-073. We thank Kevin Macaulay, Michelle Dobler, and Sydney Gallagher from the Neely lab and Hunter Noren, Samantha Buckley, and Thomas Ingalls from the NSU GIS and Spatial Ecology lab for support with field-based coral collections. We thank the University of Florida ICBR NextGen DNA Sequencing core (RRID:SCR_019152). We thank the entire Resistance Research Consortium for helpful guidance and thoughtful discussion surrounding these coral samples over the past few years.

## CRediT Roles

**Cynthia C. Becker**: Formal analysis, Writing - original draft (lead), Visualization, Data curation (lead). **Allison R. Cauvin**: Investigation, Writing – review & editing, Data curation (supporting). **Karen L. Neely**: Investigation, Conceptualization, Writing – original draft (supporting), Visualization. **Caroline Dennison**: Investigation, Writing – review & editing. **Andrew C. Baker**: Investigation, Writing – review & editing. **Brian K. Walker**: Conceptualization, Funding acquisition, Project Management, Writing – review & editing. **Julie L. Meyer**: Conceptualization, Resources, Supervision, Writing – review & editing.

## Notes

Study Funding: Florida Department of Environmental Protection Office of Resilience and Coastal Protection, Southeast Region awards B8A48D, C0CB08, C0B9A6, C1250A, and C2205D to BKW. Coral monitoring was partially supported by funding to KLN in 2019-2023 through Florida Department of Environmental Protection Coral Protection and Restoration Program (awards B40346, B54DC0, B77D91, B967BC, and C01957) and the National Fish and Wildlife Foundation in 2022-2023 (award 0302.21.071754).

### Competing Interest Statement

The authors have declared no competing interest.

