## Supplementary Figures for "Markers of resilience to stony coral tissue loss disease and probiotic potential in the microbiome of the threatened coral, *Orbicella faveolata*"

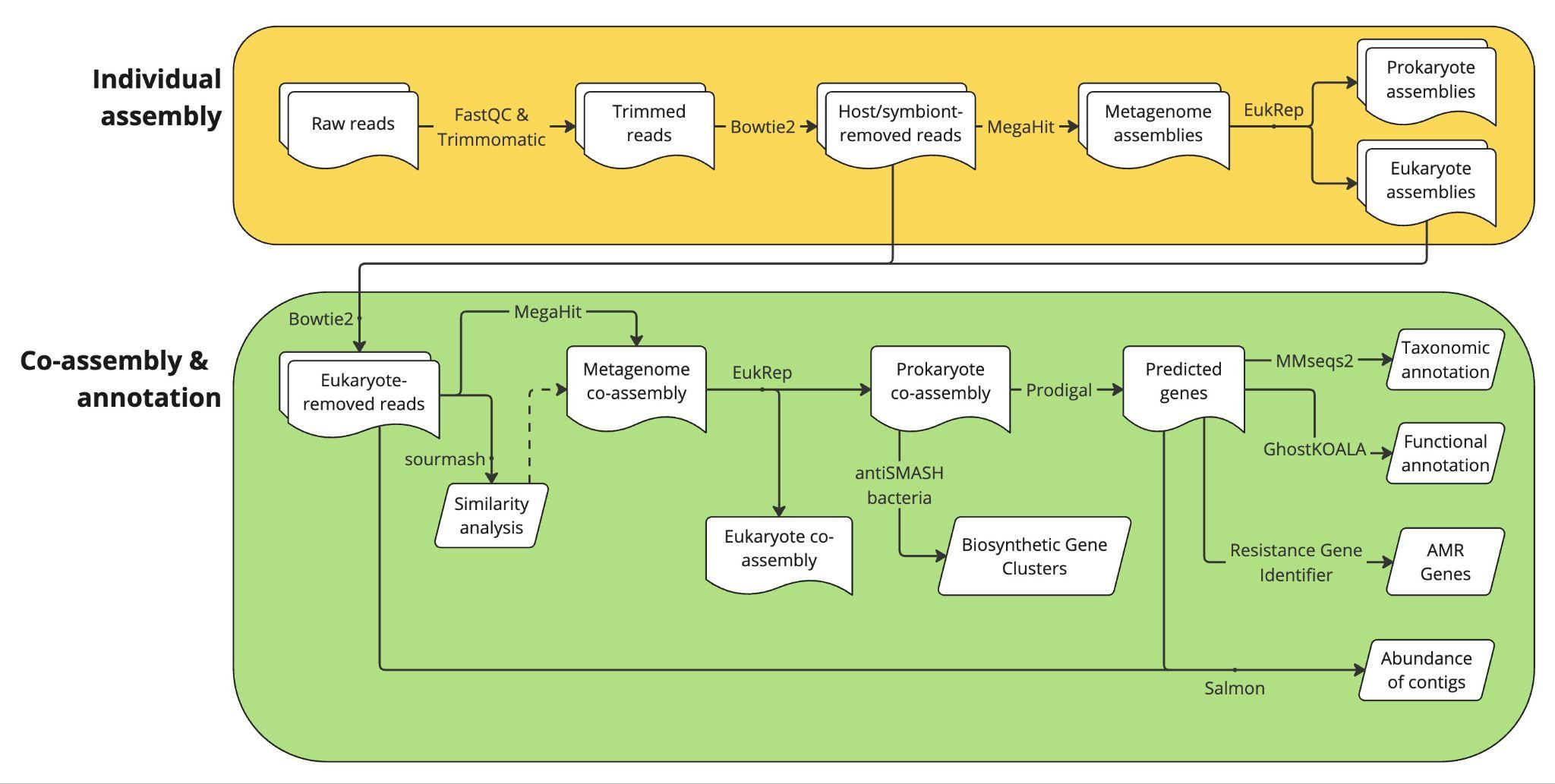

**Figure S1**. Graphical representation of the bioinformatic pipeline used to enrich prokaryotic gene content through host and eukaryotic DNA removal, followed by prokaryotic co-assembly generation and annotation.

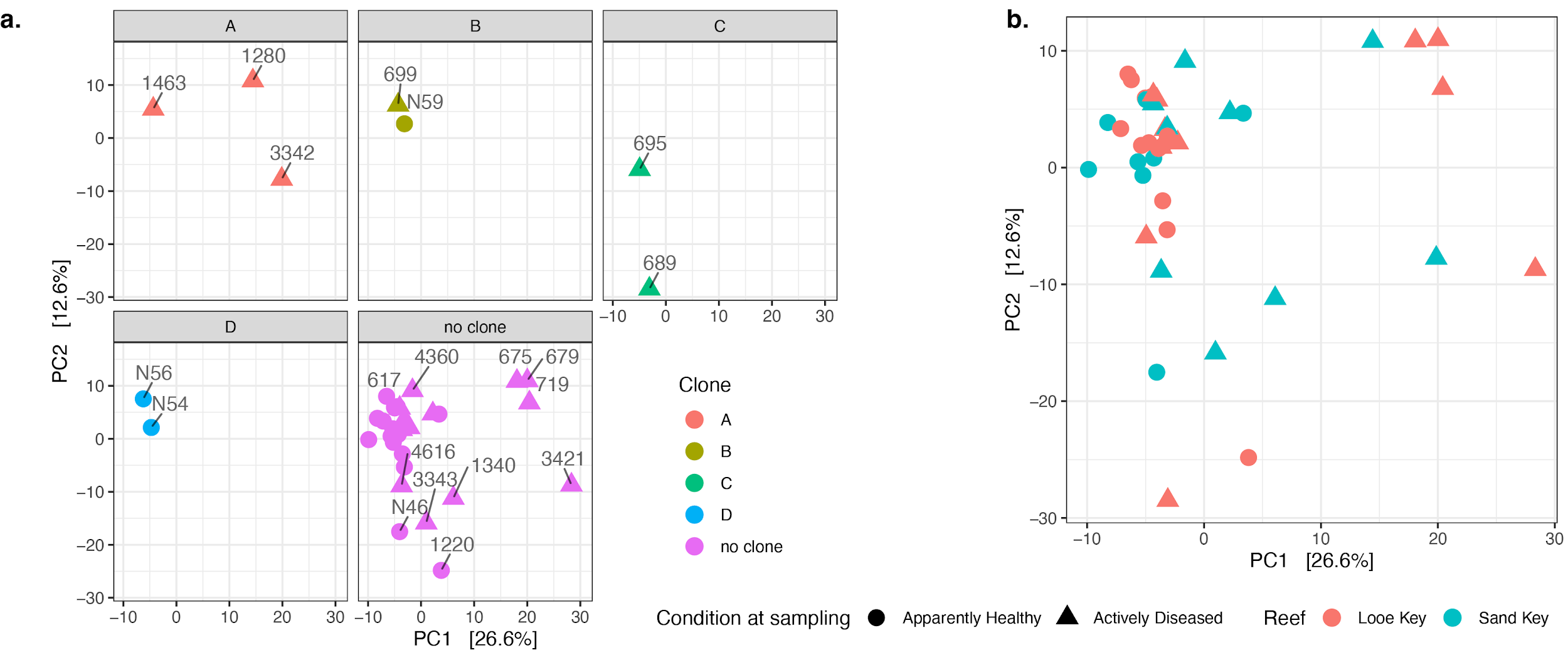

**Figure S2.** Coral genotype (defined in Klein et al. (2024)) and reef location did not explain coral prokaryotic functional microbiome beta diversity (PERMANOVA, p > 0.05). a) Principal components analysis of coral prokaryotic functional microbiome beta diversity (Aitchison distance) colored by like colony genotypes, except in the case of “no clone”, where colonies are all distinct genotypes. The genotype letters are arbitrary. The colony ID’s are labeled in panels a-d for ease of comparison across other graphs and tables. The lack of tight clustering between genotypes led us to treat genotypes as individuals (not combine the data). b) Principal components analysis as in (a) except with points colored by the reef from which the coral originated (either Looe or Sand Key) in the lower Florida Keys. The shapes in both panels represent the presence of disease on the colony at the time of sampling.
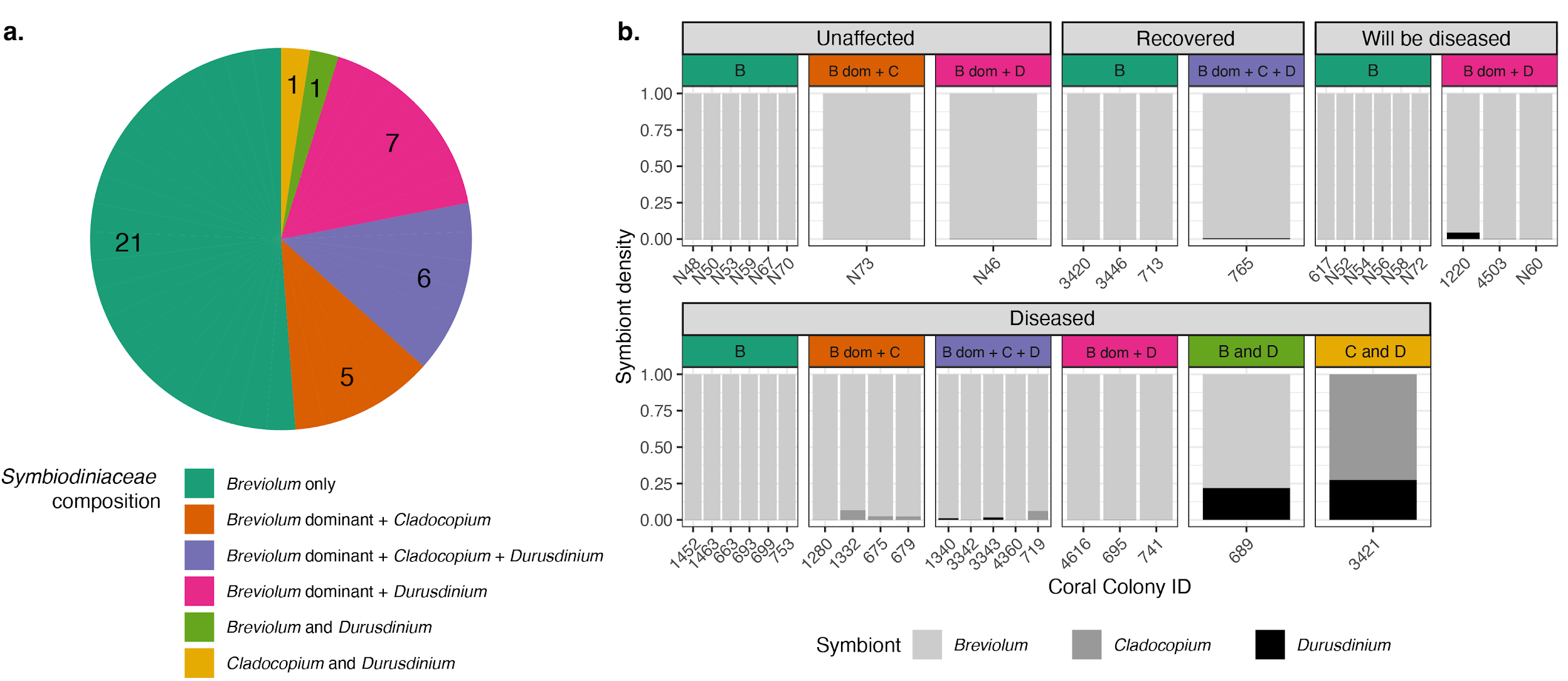

**Figure S3.** Most sampled *Orbicella faveolata* corals harbored primarily *Breviolum* photoendosymbionts. a) The proportion of the 41 sampled *O. faveolata* colonies harboring different groups of photoendosymbiont genera. Inset numbers represent the number of corals in each group. In the legend, a dominant symbiont reflects one that is >90% of the community, with the other representing <10%. In the case of the coral with *Breviolum* and *Durusdinium* and the coral with *Cladocopium* and *Durusdinium,* both symbiont genera represent more than 10% of the community. b) Proportion of different genera of *Symbiodiniaceae* photoendosymbionts in each coral. The x axis shows each coral colony ID. In some cases, the proportion of a symbiont may be too small to be visualized in the plot (e.g. *Durusdinium* density in coral colony 741). The graphs are separated based on overall colony condition at the time of sampling.

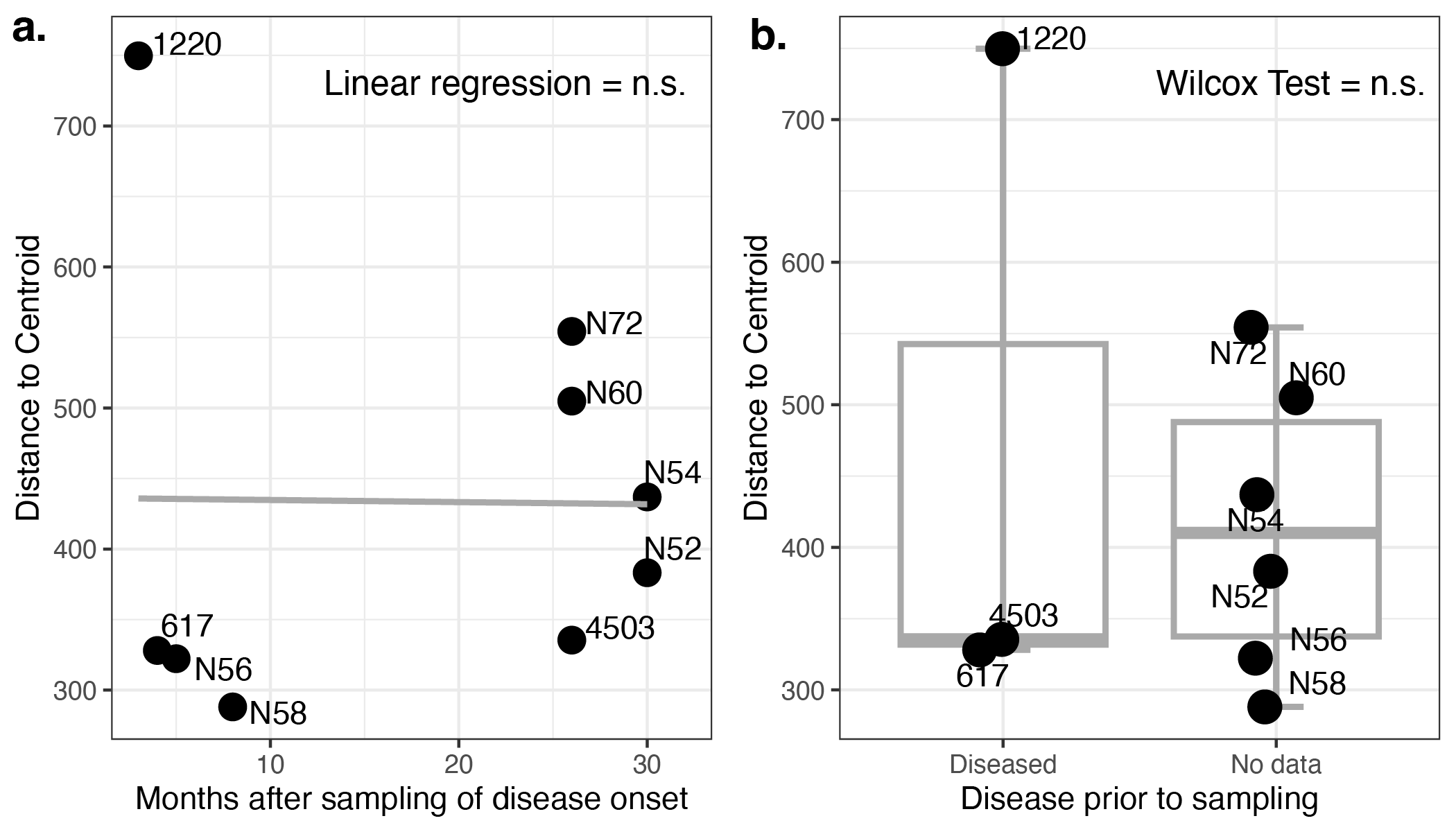

**Figure S4.** Coral functional microbial beta diversity dispersion in “will be diseased” colonies is not influenced by future onset or past history of disease (linear regression (a) p > 0.05, Wilcox test (b) p > 0.05). a) Linear regression between the number of months following sampling that disease was observed in colonies that were apparently healthy at sampling (“Will be diseased” fate). b) Box and whisker plot comparing history of disease between colonies of “will be diseased” fate. “No data” colonies can be presumed to be apparently healthy as monitoring was prevalent on reefs, but only diseased colonies were tagged prior to sampling.

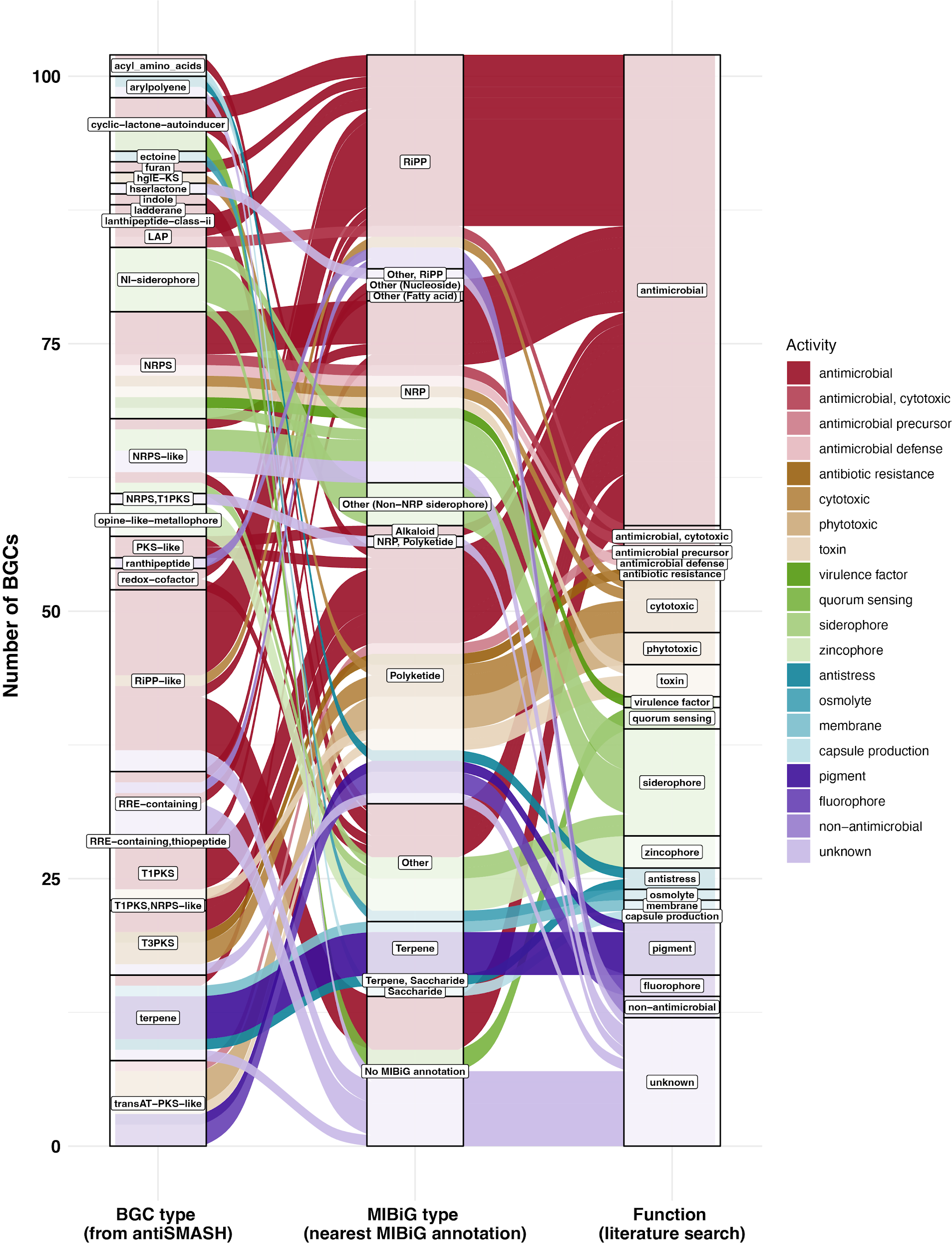

**Figure legend on next page.**

**Figure S5.** The coral microbiome coassembly contained 102 biosynthetic gene clusters (BGC’s) encoding diverse compound types. Alluvial plot displays the number of BGCs by type of BGC as returned by antiSMASH. The color of the bands reflects the likely activity or function of the BGC (third column). To infer the activity or function of the BGC, we used the closest match (if available) for the BGC to the Minimum Information about a Biosynthetic Gene cluster (MIBiG) database and reported the compound it encoded and the associated citation. For the MIBiG-generated compound, we inferred the activity or function of the compound based on the literature cited within MIBiG and related literature searches, which are reported in Table S3.

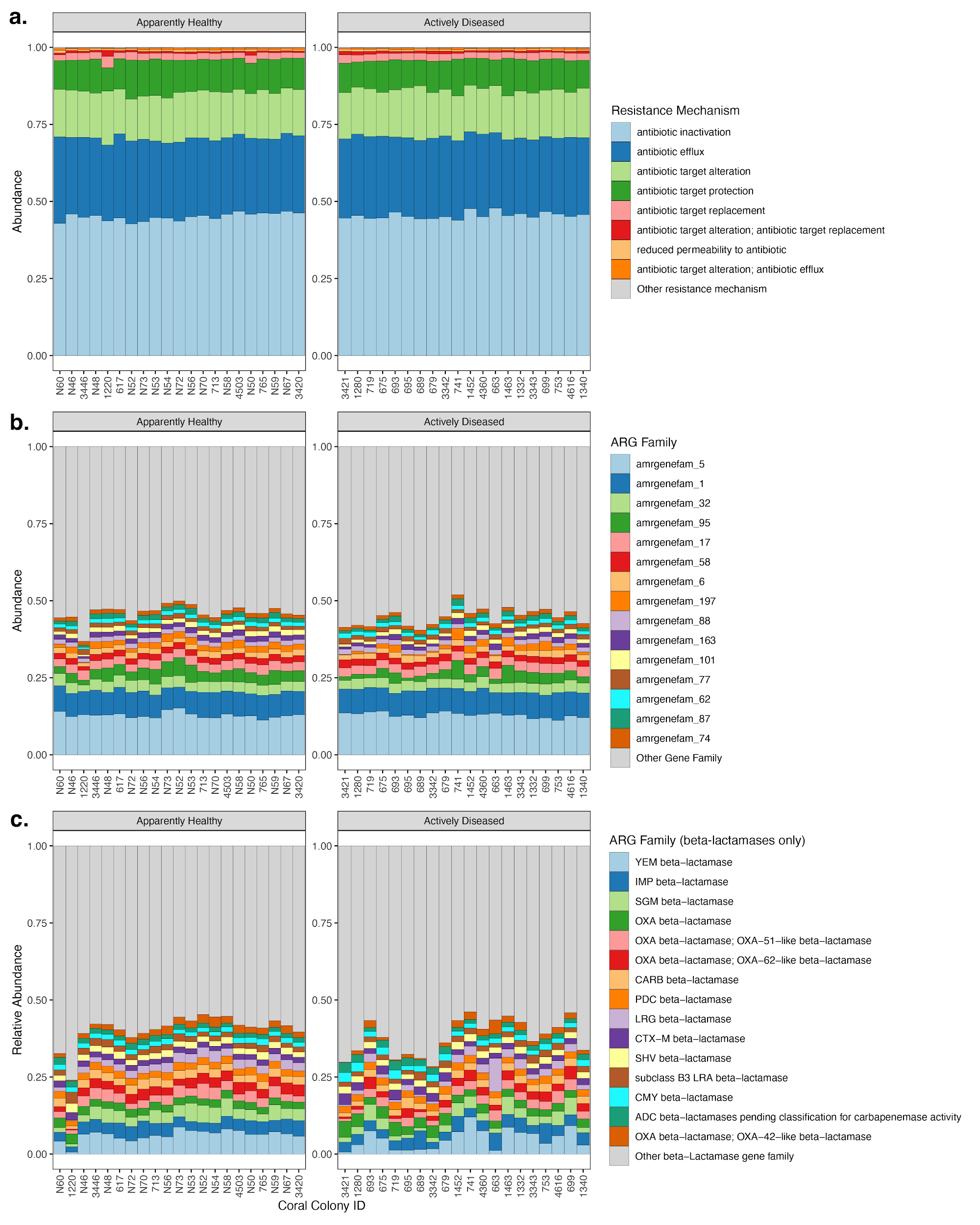

**Legend on next page**

**Figure S6**. The coral “resistome” included approximately 200,000 potential antibiotic resistance genes (ARGs) across all coral samples, including over 50,000 beta-lactamase genes. a) The ARGs fell into a variety of resistance mechanisms. The top eight resistance mechanisms are shown, and “Other resistance mechanism” includes five different mechanisms or a combination of mechanisms. b) Diverse ARG families make up the coral resistome. The top 15 most common gene families are displayed. The “Other Gene Family” category includes the remaining 422 different gene families. c) The relative abundance of the top 15 most abundant beta-lactamase genes make up less than 46% of all beta-lactamase genes. The remaining “Other beta-Lactamase gene family” category includes 238 families.

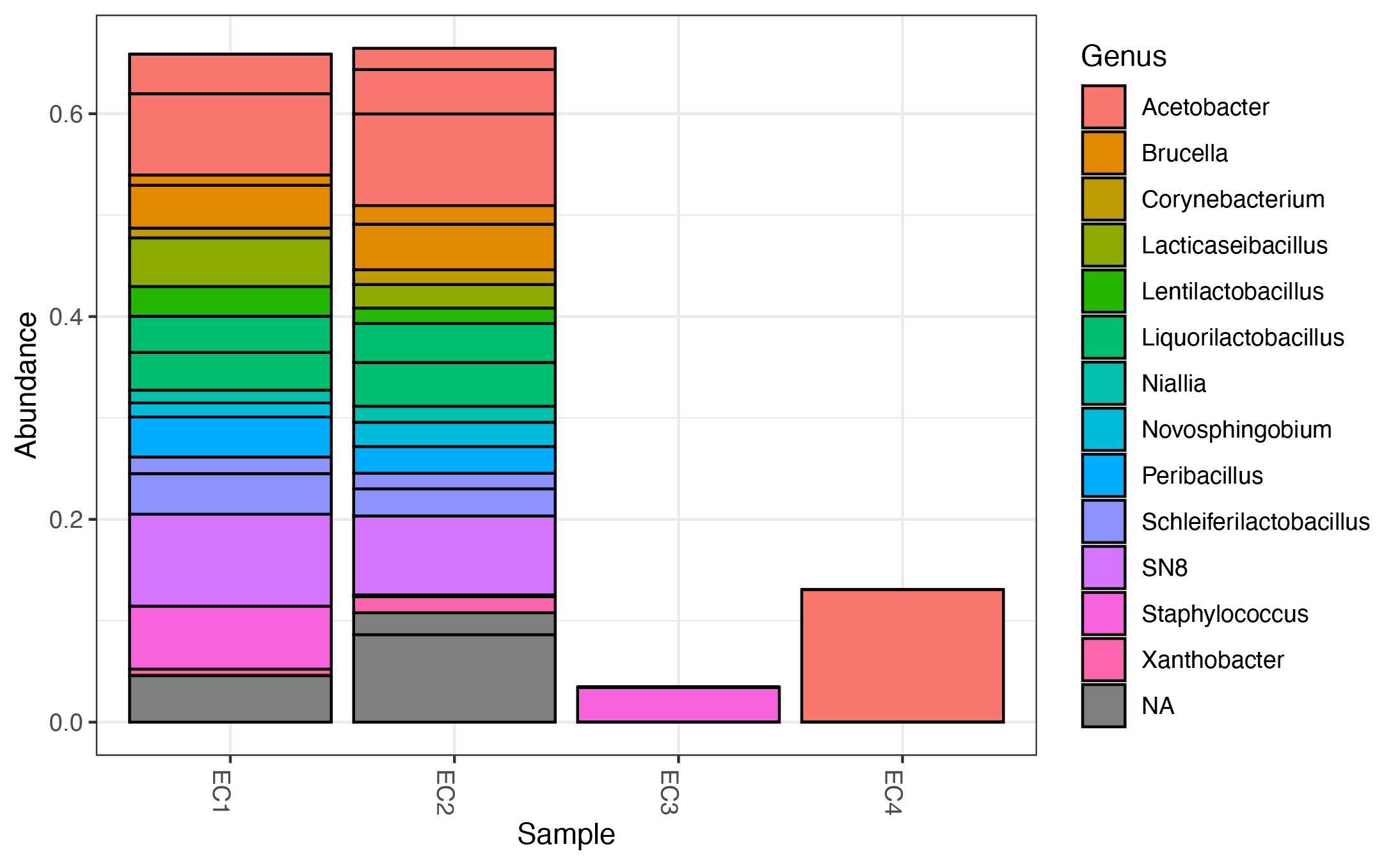

**Figure S7.** The DNA extraction blanks harbored varied genera, as determined by 16S rRNA gene sequencing. The stacked bar plot shows relative abundance of the top 20 most abundant (out of 208) ASVs across the four blank samples. A complete list of taxa and abundances of ASVs identified in the DNA extraction blanks can be found in Table S7.
