## Supplementary Information for "Markers of resilience to stony coral tissue loss disease and probiotic potential in the microbiome of the threatened coral, *Orbicella faveolata*"

### Methods

##### Study area and collection

Core samples were taken from 41 *Orbicella faveolata* colonies at two reef sites in the lower Florida Keys: Looe Key and Sand Key (Figure 1a). Both sites are spur-and-groove forereefs, and the bases of all colonies were at 3 to 9 meters depth. The two sites are approximately 49 km apart, and within each site, sampled corals were all within 0.14 km (Looe Key) to 0.3 km (Sand Key) of each other.

The selected corals were part of a long-term SCTLD intervention and monitoring project across multiple species and sites [[1–3]](https://paperpile.com/c/AriWTH/1SLWT+0nsZ5+0jgJU). Samples from the selected corals were collected by the SCTLD Resistance Research Consortium (RRC), which included the measurement of over a dozen holobiont traits [[4]](https://paperpile.com/c/AriWTH/f4Tyw) SCTLD was first documented at Looe Key in April 2018, and at Sand Key in early-mid 2019. In-water amoxicillin interventions and fate-tracking were initiated at Looe Key in March 2019 and at Sand Key in October 2020. Each site was visited approximately every two months, and any coral affected by SCTLD was tagged with a numbered cattle tag, mapped and photographed for revisitation, and treated on active disease lesions with a topical amoxicillin paste. The paste was an 8:1 by weight mixture of Ocean Alchemist’s Base2b (CoralCure) mixed with 98% amoxicillin trihydrate powder no more than 24 hours before application and placed into 60cc catheter syringes which were taken underwater. The paste was applied by hand in an approximately 1 cm wide line across any active lesions and pressed into the skeleton and coral tissue. Prior to June 2021, these efforts included 73 *O. faveolata* colonies at Sand Key and 547 *O. faveolata* colonies at Looe Key. From these candidates, we selected 16 colonies from Looe Key and 11 from Sand Key for sampling. Colonies were selected to be geographically near each other and to represent individuals who showed frequent reinfections (active lesions during most semi-monthly visits) and those who had only had active SCTLD lesions once. Additionally, we selected 15 “resistant” colonies (7 at Sand Key, 8 at Looe Key) that did not appear to have ever had SCTLD. Though we had not specifically fate-tracked these corals before selection, we chose colonies that had nearly 100% coral cover and were within the regularly monitored areas, and can thus assume they had not had any SCTLD lesions prior to selection and sampling. After sampling, all corals (including those tagged as resistant) were revisited approximately every two months until July 2024 for monitoring and treatments if necessary. During each visit (before and after sampling) the health status of the coral as well as the number of treatments (if applied) was recorded. A treatment was defined as the application of amoxicillin paste to an active lesion along the live tissue border. As such, if a single lesion affected multiple tissue isolates, that lesion could receive multiple treatments. Treatments could range in size from ~1 cm in length to >100 cm in length. Treatments were applied as needed throughout the entire colony, and were summed across all previous monitoring events (from 2019 onwards).

Cores from the 41 colonies were collected on June 12-13, 2021. Divers on SCUBA used a clean, separate 12 mm leather punch to extract a 1 cm core from apparently healthy tissue on each colony. Separate cores were collected in similar locations on the colony for metagenomics and for symbiont analyses. Core sites on each colony were selected to be distant from any active SCTLD lesions or other sources of recent mortality (Figure 1C). Each core was placed into a labeled bag and immediately transported to the surface for processing while the coring team moved to the next colony (<10 minutes from coring to processing). At the surface, tissue samples were preserved in DNA/RNA Shield (Zymo Research, Irvine, CA, USA) and kept on ice during transportation back to the lab. All samples were subsequently stored at −80ºC until processing.

##### Colony characteristics

The initial selection of *O. faveolata* colonies was based on “fully resistant” (no past or present disease lesions), “high resistance” (had disease lesions once, but after treatment never developed new lesions), and “low resistance” (repeatedly developed disease lesions). However, in the three years of fate-tracked monitoring that followed sampling, some corals shifted from one category into another, and so we developed new categories of colony fate that incorporated the full health histories of the colonies. Each colony was classed as either: 1) “diseased” (had SCTLD before sampling, during sampling, and after sampling), 2) “will be diseased” (was healthy at the time of sampling, but developed SCTLD lesions at some point afterwards – time of lesion development varied from 4 – 30 months), 3) “recovered” (coral had SCTLD before sampling, but not during sampling and at no point after sampling), or 4) “unaffected” (coral was never observed with SCTLD).

The coral host genotype was identified from adjacent samples analyzed by and reported in Klein et al [[5]](https://paperpile.com/c/AriWTH/yE1Jy). Of the 41 colonies, 33 were identified as unique genotypes, while four sets of clones were identified (three sets of two clones, and one set of three clones; Figure S2). Algal symbiont composition was determined using an actin-based quantitative PCR assay as previously described [[6]](https://paperpile.com/c/AriWTH/r6ywe).

##### Coral microbiome approach background

#### Although the coral holobiont was one of the earliest systems for shotgun metagenomic sequencing [[7, 8]](https://paperpile.com/c/AriWTH/ai1O9+Ixm3K), coral metagenomic studies that target bacterial functional genes are limited due to technical challenges. There is a stark difference in genome size between the host and symbiotic algae compared to prokaryotes. The *O. faveolata* genome is approximately 470 - 536 Mb while algal symbionts in the genus *Breviolum*, the dominant algal genera in this study, have genomes of approximately 616 Mb [[9–11]](https://paperpile.com/c/AriWTH/eoNJT+7p0J5+sFasi). Other Symbiodiniaceae genera that associate with *O. faveolata* may have even longer genomes, with *Durusdinium* and *Cladocopium* having genome sizes from 670 to over 1,000 Mb, respectively [[12, 13]](https://paperpile.com/c/AriWTH/RAzb9+CIRfU). In contrast, the average genome size of cultivated coral-associated prokaryotes is 4.2 Mb [[14]](https://paperpile.com/c/AriWTH/xMkBd). The disparity between genome sizes – specifically a difference of two to three orders of magnitude – results in a bias of metagenomic reads in favor of the coral host and Symbiodiniaceae, thereby complicating studies of the functional contributions of coral-associated prokaryotes to host health. As a result, existing metagenomic studies of SCTLD often report elevated numbers of bacterial and archaeal genes in diseased tissues, which contain a shifted microbial community and potentially a reduced signal from the tissue-associated algal symbionts [[15, 16]](https://paperpile.com/c/AriWTH/0FjK6+N2kGU). To focus on the prokaryotic component of apparently healthy coral, some studies use additional filtration of tissues to enrich microorganisms [[17, 18]](https://paperpile.com/c/AriWTH/XnMKR+NEN44). Alternatively, sequencing deeply to recover a large proportion of microbial reads is essential when host contamination is expected to be above 90% [[19]](https://paperpile.com/c/AriWTH/KJ6Eo). Since we are interested in signatures of resilience and resistance to disease, targeting apparently healthy tissue before it is actively degraded is key. Additionally, using deep sequencing enables the recovery of the prokaryotic component of the coral holobiont.

##### DNA Extraction and Library preparation

Samples were randomized into batches of eight for DNA extraction. For DNA extraction, coral cores were thawed on ice. Once thawed, the coral tissue was scraped from cores using flame-sterilized scalpels and forceps over a sterile Petri dish. The scraping involved collecting a portion of the coral surface mucus as well as 1-2 polyps into sterile tubes containing 1mL of phosphate-buffered saline with ethylenediaminetetraacetic acid. A sterile micropestle was then used to mechanically break up the tissue and create a slurry. The resulting tissue slurry was then added to the Qiagen QIAamp DNA Microbiome Kit (Qiagen, Germantown, MD, USA) to preferentially extract the microbial DNA instead of host DNA according to the manufacturer’s protocol. Reagent blanks were generated for each batch of DNA extractions wherein the blanks were processed alongside experimental samples, without the addition of coral biomass. Nucleic acid extracts were quantified with a DeNovix high-sensitivity dsDNA fluorescence assay to verify extraction success.

The extracted DNA was then submitted to the University of Florida’s Interdisciplinary Center for Biotechnology Research NextGen DNA Sequencing core (RRID:SCR_019152) for library preparation and sequencing. The NEBNext® Ultra™ II DNA Library Prep Kit for Illumina (New England Biolabs, Ipswich, MA, USA) was used to prepare the sample libraries with the NEBNext® Multiplex Oligos for Illumina® (96 Unique Dual Index Primer Pairs; New England Biolabs, Ipswich, MA, USA). Prepared sample libraries were then sequenced over four lanes on an Illumina NovaSeq 6000 (Illumina, San Diego, CA, USA) in a 150-bp paired-end format. Raw fastq files are openly available through the NCBI Sequence Read Archive under accession number PRJNA925892.

##### Reagent Blank Analysis

Due to low biomass, we did not sequence the reagent blank samples with the metagenomic sequencing on the Illumina NovaSeq. Instead, we subjected the four blanks to 16S rRNA gene sequencing. The V4 hypervariable region of bacterial and archaeal 16S rRNA genes was amplified with the Earth Microbiome Protocol [[20]](https://paperpile.com/c/AriWTH/yn3Ep) using the 515F [[21]](https://paperpile.com/c/AriWTH/ryPVN) and 806R [[22]](https://paperpile.com/c/AriWTH/g5ZTe) primers, as previously described [[23]](https://paperpile.com/c/AriWTH/UEZrG). Amplicon libraries were submitted to the University of Florida’s Interdisciplinary Center for Biotechnology Research NextGen DNA Sequencing core (RRID:SCR_019152) for sequencing on an Illumina MiSeq with the 2 x 150 base pair (bp) v2 cycle format. Following sequencing, we used cutadapt v3.4 [[24]](https://paperpile.com/c/AriWTH/8tsgp) followed by dada2 (v1.30.0) to parse the fastq files into amplicon sequence variants (ASV) [[25]](https://paperpile.com/c/AriWTH/Rb5I4). We used the following parameters for filterAndTrim: truncLen=c(145,145), maxN=0, maxEE=c(2,2), truncQ=2, and rm.phix=TRUE. Following filtering, learning error rates, and merging of paired reads, we removed chimeric sequences and taxonomically identified each ASV to the species level with the SILVA database v.138.2 [[26]](https://paperpile.com/c/AriWTH/TKHPC). The resultant table of ASV counts and taxonomy are available in Table S7.

##### Bioinformatics

An overview of the bioinformatics pipeline we employed to target bacteria and archaea within the coral holobiont is outlined in Figure S1. We assessed the quality of raw reads using FastQC v.0.11.7, then filtered the raw reads using trimmomatic v.0.39 (Bolger, Lohse, and Usadel 2014). The parameters included LEADING:3; TRAILING:3; SLIDINGWINDOW:4:20; MINLEN:50. To remove host and endosymbiont DNA and begin enriching for prokaryotic genomic content, we used Bowtie2 v.2.4.2 with parameters: --un-conc-gz to retain unaligned reads [[27]](https://paperpile.com/c/AriWTH/FSICV) to map trimmed reads against the following genomes: the *O. faveolata* genome (NCBI Assembly Accession ID GCA_001896105.1), *Breviolum minutum* (NCBI Assembly Accession ID GCA_000507305.1; [[11]](https://paperpile.com/c/AriWTH/sFasi)), *Symbiodinium* sp. Clade C Y103 genome (*Cladocopium*; NCBI Assembly Accession ID: GCA_003297045.1; [[28]](https://paperpile.com/c/AriWTH/jsieO)), and *Durusdinium sp.* (NCBI Project ID: [PRJDB10306](https://www.ncbi.nlm.nih.gov/bioproject/?term=prjdb10306)). The number of reads retained after this initial quality filtering are listed in Table S2. We assembled the host/endosymbiont-cleaned reads from each of the 41 samples individually into contigs using MegaHit v.1.1.3, employing the –metalarge parameter that is optimized for assembling large, complex datasets [[29]](https://paperpile.com/c/AriWTH/aehlF). This resulted in a metagenome assembly for each *O. faveolata* colony*.* These metagenome assemblies are available at Zenodo DOI 10.5281/zenodo.11493775 [[30]](https://paperpile.com/c/AriWTH/W15bp). These metagenome assemblies were further assessed using QUAST v.5.2.0 [[31]](https://paperpile.com/c/AriWTH/AT7aB) to determine statistics such as total length, largest contig, and number of contigs (Table S2).

To evaluate the broad prokaryotic and eukaryotic content of the metagenome assemblies, we used EukRep v.0.6.7 [[32]](https://paperpile.com/c/AriWTH/pgTw1) with the parameters --min 1000 and --prokarya to keep only contigs over 1000 bp and to parse prokaryotic versus eukaryotic contigs. We then counted the number of contigs in each group and used QUAST v.5.2.0 using default parameters to generate summary statistics on the prokaryotic portion of the metagenome assemblies (Table S2).

Following host and symbiont removal, we noted that prokaryotic reads were only 9.66 ± 4.31% of the total assembly length, so we further enriched the prokaryotic component of the host/endosymbiont-cleaned reads prior to generating a co-assembly (Table S2). We used Bowtie2 v.2.4.2 [[27]](https://paperpile.com/c/AriWTH/FSICV) to map the host/endosymbiont-cleaned reads to the eukaryotic portion of the metagenome assembly generated by EukRep. We first used bowtie2-build to create an index of the eukaryotic portion of the metagenome assembly using default parameters. Then, we used bowtie2 to map the reads to the eukaryotic indices and retain reads that did not map with the flags --un-conc-gz, --phred33, and -q. which we refer to as eukaryote-removed reads.

In order to recover longer contigs for greater bacterial functional annotation, maintain rare taxa, and to enable cross-sample comparison, we generated a co-assembly of all 41 coral samples. Prior to generating a co-assembly, we used sourmash v.4.5.0 [[33]](https://paperpile.com/c/AriWTH/0H9nS) to generate hash sketches and investigate the similarity of eukaryote-removed reads across the samples to verify samples would co-assemble well. Specifically, we used sourmash sketch dna (parameters k=31), followed by sourmash compare, followed by sourmash plot (parameters --labels). Since the hash sketches did not differ by reef (Looe Key versus Sand Key), we concluded the genomic content was similar and combined them into a co-assembly. To generate the co-assembly, we used MegaHit v.1.1.3 [[29]](https://paperpile.com/c/AriWTH/aehlF) with parameters --min-contig-len 1000, -t 100, --presets meta-large. The meta-large preset is for large and complex metagenomes, which we expected for coral metagenomes. We evaluated the resultant assembly with QUAST v.5.2.0 using default parameters. Despite multiple steps of eukaryotic read removal, we anticipated there might still be eukaryotic content. Therefore we used EukRep v.0.6.7 again with the same parameters as before to assess the relative proportion of the co-assembly that was prokaryotic and eukaryotic. We used the prokaryotic portion of the co-assembly reflecting 41 *O. faveolata* corals for subsequent annotation and analysis.

To predict protein-coding genes, we used Prodigal v.2.6.3 [[34]](https://paperpile.com/c/AriWTH/FUNSw) with the parameters “-f gff” to output a gff file and “-p meta” because the input file was a metagenome. We additionally generated a nucleotide fasta of the predicted protein coding genes (-d flag) for downstream abundance calculation. We annotated the predicted genes (amino acid fasta) with MMseqs2 v.2/14, parameters --search-type 1, --tax-lineage 1, --threads 50, --compressed 1), employing the GTDB database v.214.1 to generate taxonomic annotations [[35]](https://paperpile.com/c/AriWTH/MX7Tw). For functional annotations, we used the KEGG “Prokaryotes + Viruses” database using the web-based GhostKOALA tool to annotate the prokaryote predicted genes (amino acid fasta) [[36]](https://paperpile.com/c/AriWTH/8ntSQ). We quantified the abundance of predicted genes from the prokaryote co-assembly using Salmon v.1.10.1 [[37]](https://paperpile.com/c/AriWTH/AGggx). We first indexed the nucleotide fasta of the predicted genes (salmon index, parameters -k 31), then mapped the eukaryote-removed reads to the index using salmon quant (parameters --meta, --libType A, -p 36). We saved the “NumReads” column as integers from the Salmon output for each coral sample and aggregated it into one abundance table. The resultant table represented read counts for each predicted gene in the prokaryotic co-assembly for each coral sample, and was used for downstream analyses. The predicted proteins, annotations, and abundance table are publicly available at Zenodo DOI 10.5281/zenodo.11493775 [[30]](https://paperpile.com/c/AriWTH/W15bp).

We used the resistance gene identifier (RGI) v.6.0.2 to annotate the predicted genes from the prokaryote co-assembly (amino acid fasta) using the Comprehensive Antibiotic Resistance Database (CARD) database, with the parameters: --clean --include_loose -t protein -n 20 [[38]](https://paperpile.com/c/AriWTH/BXL9F). We used the “--include_loose” parameter to include “loose” hits to the database in addition to strict and perfect hits. This parameter is better for detecting new or distant homologs of antimicrobial resistance genes, and since this coral microbiome dataset is environmental, we expected this parameter to be better suited for our dataset, as we would not expect many clinically-relevant antimicrobial resistance genes. To identify BGCs, we used antiSMASH v.7.0.0 on the prokaryotic co-assembly generated by EukRep (nucleotide fasta containing contigs), using the parameters: --taxon bacteria --tigrfam --pfam2go --cc-mibig --rre --genefinding-tool prodigal-m --cpus 25 [[39]](https://paperpile.com/c/AriWTH/Nm2Pq).

##### Statistical analyses

For statistical tests, we used the read counts (“NumReads” from Salmon as integers) of each predicted gene across coral samples (247,254 genes). We analyzed the functional microbial beta diversity using Aitchison distance to account for the compositional structure of the sequencing data [[40]](https://paperpile.com/c/AriWTH/48oSx). To do this we applied a centered log-ratio (CLR) transformation on the abundance-filtered read counts using the microbiome package v.1.24.0 in R v.4.3.2 [[41]](https://paperpile.com/c/AriWTH/PaSI2), which incorporates a pseudocount of min(relative abundance)/2 to relative abundances of zero prior to conducting the CLR transformation. We then conducted the ordination and principal components analysis which uses Euclidean distance with phyloseq v.1.46.0 [[42]](https://paperpile.com/c/AriWTH/lnzqv). To test our hypotheses and examine the extent to which colony-specific covariates (coral genotype, fate, disease condition at sampling, Symbiodiniaceae composition, and reef location) explained variability in the functional microbiome, we used the adonis2 function [[43]](https://paperpile.com/c/AriWTH/4Q7GU) in vegan v.2.6.6.1 to conduct a PERMANOVA. We first investigated coral genotype-based changes to the functional microbiome to justify treating coral genotypic clones [[5]](https://paperpile.com/c/AriWTH/yE1Jy) as individuals or aggregating their functional microbiomes for further analyses. Based on the results, we chose to treat the genotypic clones as separate individuals for all subsequent analyses (Figure S2a). To investigate how dispersion of the prokaryotic microbiome changed with colony fate, condition at sampling, or Symbiodiniaceae community composition, we calculated the distance to centroid in the principal components analysis using the betadisper function in vegan v.2.6.6.1. We hypothesized dispersion would increase in diseased colonies [[44]](https://paperpile.com/c/AriWTH/LhOzZ). Then, based on the normal distribution of the distances, we conducted a Kruskal-Wallis or ANOVA test followed by either a Wilcoxon or Tukey post-hoc test to investigate differences across groupings.

To identify genes that may be involved in disease resistance, we conducted a differential abundance test in corncob between unaffected corals (baseline) and the other colony fates: recovered, will be diseased, and diseased. To glean biological information from the differential abundance analysis, we conducted the differential abundance test only on the 31,695 genes annotated by ghostKOALA. We used corncob v.0.4.1 [[45]](https://paperpile.com/c/AriWTH/lioZk) to test the hypotheses that microbial functional genes differed in abundance between apparently healthy tissue in corals from unaffected fates and either recovered, will be diseased, or diseased coral fates. We controlled for the composition of Symbiodiniaceae as that significantly structured the functional microbial community. Corncob modeled relative abundances from the input count data, and differential abundance was modeled as a linear function of coral fate. Significant differences were evaluated using the parametric Wald test, with a Benjamini-Hochberg false discovery rate correction of 0.05 to account for multiple comparisons. Coefficients including 95% confidence intervals were plotted to display differentially abundant genes. To plot the data, the KEGG annotations were retrieved for each KO ID using the KEGG API in R and parsed into different levels.

To test whether repeated amoxicillin treatments impacted the richness and diversity of beta-lactamase genes, we subset genes with the string “beta-lactamase” in the AMR Gene Family annotation, and used the package breakaway v.4.8.4 to estimate the richness of beta-lactamase genes within each *O. faveolata* colony [[46]](https://paperpile.com/c/AriWTH/oJuEz). We then tested for the effect of amoxicillin treatments and recent treatment history on estimated beta-lactamase gene richness using the betta function in the breakaway package [[47]](https://paperpile.com/c/AriWTH/xmHtz). We additionally tested for the effect of these variables on the beta diversity of beta-lactamase genes by generating a PCA and PERMANOVA test using methods described above. The variables we tested for their effect on richness and beta diversity included both raw numbers of total amoxicillin treatments prior to sampling (continuous variable) and the number of treatments of amoxicillin scaled to the estimated live tissue area in meters squared (continuous). Additionally, to test whether treatment history affected diversity of beta-lactamases, we used a categorical variable indicating a colony was never treated, treated more than two months prior to sampling, or treated within two months prior to sampling.

We inferred the potential activity or function of each BGC by using matches to the Minimum Information about a Biosynthetic Gene cluster (MIBiG) database. AntiSMASH reported matches of any portion of a BGC to entries within the MIBiG database. We chose the match with the highest similarity score and reported the MIBiG accession number, similarity score, compound, source organism, and relevant citation from the MIBiG database in Table S3. We used experimental data and information from the MIBiG reference or in some cases additional literature searches to identify the activity or function of the reported compound. The similarity scores were low for the majority of MIBiG matches, which we expected given the complex nature of coral and environmental samples. Therefore, it is likely that the close matches from the coral holobiont may not be the exact compounds referenced in MIBiG. Instead, we used these matches to infer the potential function or activity (siderophore, antimicrobial, toxin, etc) for the BGC. To track how each BGC was annotated and assigned the inferred activity or function, we generated an alluvial plot.

To test whether BGCs with roles related to coral disease resilience were differentially prevalent across colony fates, we used a Fisher’s Exact Test incorporating a Monte-Carlo simulation to generate a p-value, followed by a Benjamini-Hochberg multiple test correction in R. Following manual curation of antiSMASH and MIBiG results, we used a subset of the BGCs which we hypothesized would play a role in coral disease resilience, including gene clusters annotated as “antimicrobial”, “antimicrobial, cytotoxic”, “antimicrobial precursor”, “antimicrobial defense”, “antibiotic resistance”, “cytotoxic”, “phytotoxic”, “toxin”, or “virulence factor”. CoverM v.0.6.1 was used to measure mean coverage using coverm contig with the parameters “--methods mean” and “--min-covered-fraction 10” from the eukaryote removed reads [[48]](https://paperpile.com/c/AriWTH/Ttk72) and then converted coverage to presence/absence for statistical analysis.

### Results and Discussion

##### Antibiotic Resistance Gene Diversity

The coral microbiome encoded over 200,000 ARGs, of which over 50,000 were beta-lactamase genes (Figure S6, Table S7). These ARGs were present across *O. faveolata* corals. Most ARGs encoded efflux or inactivation mechanisms of resistance, along with 13 other mechanisms of resistance (Figure S6a). These genes fell into 437 different ARG families, of which the top 15 comprised between 37 - 52% of the abundance of all ARGs (Figure S6b). Within the beta-lactamases, the most abundant families were YEM, IMP, SGM, and OXA beta-lactamases, and the top 15 beta-lactamase gene families made 24 - 46% of all beta-lactamase gene counts in colonies (Figure S6c).

##### Blank Analysis

We generated 16S rRNA gene-based analyses of four DNA extraction reagent blanks to identify potential contaminants. Of the four blanks, two contained about 17,000 reads, while the other two contained about 1,200 reads (Figure S7, Table S7). The most abundant reagent blank sequences were assigned to the following genera: *Acetobacter, Brucella, Corynebacterium, Lacticaseibacillus, Lentilactobacilus, Liquorilactobacillus, Niallia, Novosphingobium, Peribacillus, Scheiferilactobacillus, Staphylococcus, and Zanthobacter* (Figure S7). A complete list of taxa in the blanks is in Table S7.

One challenge in this study was the lack of reagent blanks that matched the full processing steps of the metagenome samples. Reagent blanks from the DNA extraction process were subjected to PCR amplification of the 16S rRNA gene for taxonomic microbiome analysis, which resulted in 208 ASVs, that substituted for reagent blanks that included shotgun metagenomic library preparation. We reasoned that library preparation would be unsuccessful with the low- to no-biomass reagent blanks and would be rejected by the sequencing center. In hindsight, this could have been overcome with the addition of mock community DNA to reagent blanks prior to sequencing, and this is a tactic recommended for future studies. We found some taxa in the results that overlapped with those found in the DNA blanks. For example, we identified one *Acetobacter* and *Corynebacterium* gene each within our differentially abundant analysis, and these were also in three or two out of the four DNA blanks, respectively. While *Acetobacter* is a genus of acetic acid bacteria associated with fuel tank corrosion, it has also been isolated from soft corals [[49, 50]](https://paperpile.com/c/AriWTH/WUFUI+26Wsy). Similarly, *Corynebacterium* has been isolated from a hard *Fungia* coral, as well as associated with human skin infections [[51, 52]](https://paperpile.com/c/AriWTH/7pTd1+pIyI7). Of the top 20 most abundant taxa within the DNA blanks, *Staphylococcus* was represented most within the differentially abundant functional genes. *Staphylococcus epidermis* is a common skin bacterium and pathogen [[53]](https://paperpile.com/c/AriWTH/HRrR6). *Staphylococcus* is also a coral opportunistic pathogen and is commonly found in the coral microbiome [[54–56]](https://paperpile.com/c/AriWTH/hCT48+qDLoA+I2YbM). While some overlap exists, there were also some taxa reassuringly absent from the DNA blanks, such as *Cutibacterium* (formerly *Propionibacterium*). Many contigs classified as *Cutibacterium,* and one BGC with a perfect similarity score matched a cutimycin gene cluster from *Cutibacterium acnes.* While *Cutibacterium acnes* is a skin commensal and opportunistic pathogen, it has also been found in coral metagenome-assembled genomes, and even been considered a potential endosymbiont [[17, 57, 58]](https://paperpile.com/c/AriWTH/UFEC5+XnMKR+ljuq8). Overall, while there was some overlap between the taxa recovered in the DNA blanks, we did not remove them from analysis given the different methodological approaches and associated bias (i.e. PCR amplification with 16S rRNA methods) as well as due to prior research linking similar genera with marine and coral habitats. In addition, sequencing errors resulting in sample “cross-talk” where sequence reads are misbarcoded is also a potential issue with the Illumina sequencing platform [[59]](https://paperpile.com/c/AriWTH/clIjP), thus taxa detected in the reagent blanks may actually be sourced from true samples. In the future, incorporating both mock community DNA and reagent blanks spiked with mock community DNA in shotgun metagenome sequencing efforts will improve our assessment of potential contamination from cross-talk, kit reagents, or users [[59]](https://paperpile.com/c/AriWTH/clIjP).
